# Arginine 65 methylation of Neurogenin 3 by PRMT1 is a prerequisite for development of hESCs into pancreatic endocrine cells

**DOI:** 10.1101/2022.03.15.484450

**Authors:** Gahyang Cho, Kwangbeom Hyun, Jieun Choi, Eunji Shin, Bumsoo Kim, Jaehoon Kim, Hail Kim, Yong-Mahn Han

## Abstract

In the developing pancreas, Neurogenin 3 (NGN3) is a key transcription factor in the cell fate determination of endocrine progenitors (EPs). Although the activation and stability of NGN3 are regulated by phosphorylation, the role of arginine methylation of NGN3 is poorly understood. Here, we report arginine 65 methylation of NGN3 is absolutely required for the pancreatic lineage development of human embryonic stem cells (hESCs) *in vitro*. First, we found inducible protein arginine methyltransferase 1 (PRMT1)-knockout (P-iKO) hESCs did not differentiate from EPs to endocrine cells (ECs) in the presence of doxycycline. Loss of PRMT1 caused an accumulation of NGN3 in the cytoplasm of EPs and blocked NGN3’s transcriptional activity in PRMT1-KO EPs. We also found that PRMT1 specifically methylates NGN3 arginine 65, and this modification is a prerequisite for ubiquitin-mediated NGN3 degradation. Our findings indicate arginine 65 methylation of NGN3 is a key molecular switch in hESCs *in vitro* permitting the differentiation into pancreatic endocrine lineages.

## Introduction

The endocrine system of pancreas—a small proportion of the pancreas as a whole— comprises small globular clusters of specialized cells, such as the Islets of Langerhans. In mammalian embryos, the cells of the pancreatic endocrine lineage develop through a series of successive stages, including definitive endoderm (DE), pancreatic endoderm (PE), endocrine progenitor (EP), and finally endocrine cells (EC) (D’Amour et al., 2005; Pan and Brissova, 2014). As PE cells arise from DE cells, they begin to exhibit strong expression of pancreatic and duodenal homeobox 1 (PDX1) at E8.5 in mice and 30 d post-conception (dpc) in humans (Martin F. Offield, Tom L. Jetton, Patricia A. Labosky, Michael Ray and Mark A. Magnuson, Brigid L. M. Hogan, 1996). PDX1-positive endodermal cells then differentiate into EPs, which exhibit transient activation of the transcription factor Neurogenin 3 (NGN3) (Jennings et al., 2020; Schwitzgebel et al., 2000; Smith et al., 2004). NGN3 is a critical determinant of EC fate that activates EC-associated genes (Guoqiang Gu, 2002; Jennings et al., 2015). NGN3^NULL^ mice fail to develop any pancreatic EC lineages (Gradwohl et al., 2000). Normally, the strong NGN3 expression of trunk EP cells falls abruptly within a few days (Apelqvist et al., 1999). Although it is clear that multi-phosphorylation of NGN3 facilitates both its ubiquitin-mediated degradation and its transcriptional activity (Azzarelli et al., 2017; Krentz et al., 2017), it is unknown whether other more elusive modifications important for NGN3 degradation or stability are involved in pancreatic development.

Protein arginine methyltransferase 1 (PRMT1), the predominant asymmetric arginine methyltransferase in mammalian cells, methylates arginine residues in conserved glycine and arginine-rich (GAR) regions of histone and non-histone proteins (Bedford and Richard, 2005; Tang et al., 2000). Through its role in arginine methylation, PRMT1 contributes to diverse cellular processes, such as epithelial-mesenchymal transition (EMT), cell cycle, DNA repair, and RNA processing (Avasarala et al., 2015; Bedford and Clarke, 2009; Gary and Clarke, 1998; Raposo and Piller, 2018). It also functions as a transcriptional coactivator via its role in forming asymmetrical dimethyl-arginines on histone 4 (H4R3me2a) (Strahl et al., 2001; Wang et al., 2001). PRMT1-mediated arginine methylation plays an important role in the stability and localization of non-histone proteins, such as FOXO1, TSC2, and progesterone receptor (Gen et al., 2020; Malbeteau et al., 2020; Yamagata et al., 2008). Recently, Prmt1- knockout (PKO) mouse embryos showed slower degradation of NGN3 from E14.5, leading to severe postnatal pancreatic hypoplasia (Lee et al., 2019). This suggests PRMT1 is associated with NGN3 stability in pancreatic development. Nonetheless, the mechanism by which PRMT1 affects pancreatic development via its regulation of the stability and activity of NGN3 is poorly understood.

To explore the importance of NGN3 arginine methylation in pancreatic development, we generated a PRMT1-inducible knockout (P-iKO) H1 hESC line. We found that although PRMT1-KO (P-KO) EPs show similar NGN3 transcriptional activity compared to WT EPs, they accumulate more NGN3 protein. Unexpectedly, this unmethylated NGN3 that accumulated because of the loss of PRMT1 did not support pancreatic EC development. In addition, we found NGN3 arginine 65 can only be methylated by PRMT1. Together, our results indicate PRMT1 plays a critical role in pancreatic EC development via its methylation of NGN3 arginine 65.

## Results

### Generating a P-iKO hESC line

To understand the role of arginine methylation in pancreatic endocrine cell development, we generated a PRMT1-inducible knockout (P-iKO) H1 hESC line (Figure S1). To do this, we inserted the pAAV-Neo_Cas9 and pAAV-Puro_siKO vectors into the AAVS1 locus in H1 hESCs using zinc-finger nucleases (ZFN-L and ZFN-R). One AAVS1 allele was targeted for constitutive Cas9 expression and the other for tetracycline-dependent (TetR) sgRNA transcription (Figure 1A, left panel). We designed the sgRNA to target human *PRMT1* exon 7, which is in the middle of the SAM-dependent methyl-transferase domain (Figure 1A, right panel). Only one targeted clone survived antibiotic selection. In normal media, P-iKO cells maintain an intact *PRMT1* genomic locus because they only produce Cas9 protein. The P- iKO cells begin expressing the sgRNA when exposed to doxycycline (dox). Then, the Cas9- sgRNA complex makes indels in the *PRMT1* locus of P-iKO cells (Figure 1B, arrows: 360, 440bp). In the presence of dox, all P-iKO hESCs died within 48 h, whereas H1 hESCs survived (Figure 1C). As expected, *PRMT1* transcripts were significantly reduced in dox-treated P-iKO hESCs compared to untreated ones (Figure 1D). Like H1 hESCs, however, P- iKO hESCs exhibited normal expression of pluripotent markers, including SOX2, TRA 1-60, OCT4, and TRA 1-81 (Figure 1E). They also showed normal expression of PRMT1 and normal methylation of histone 4 (H4R3me2a) (Figure 1F). Thus, we generated pluripotent P- iKO hESCs with an inducible P-KO system *in vitro*.

**Figure 1.**
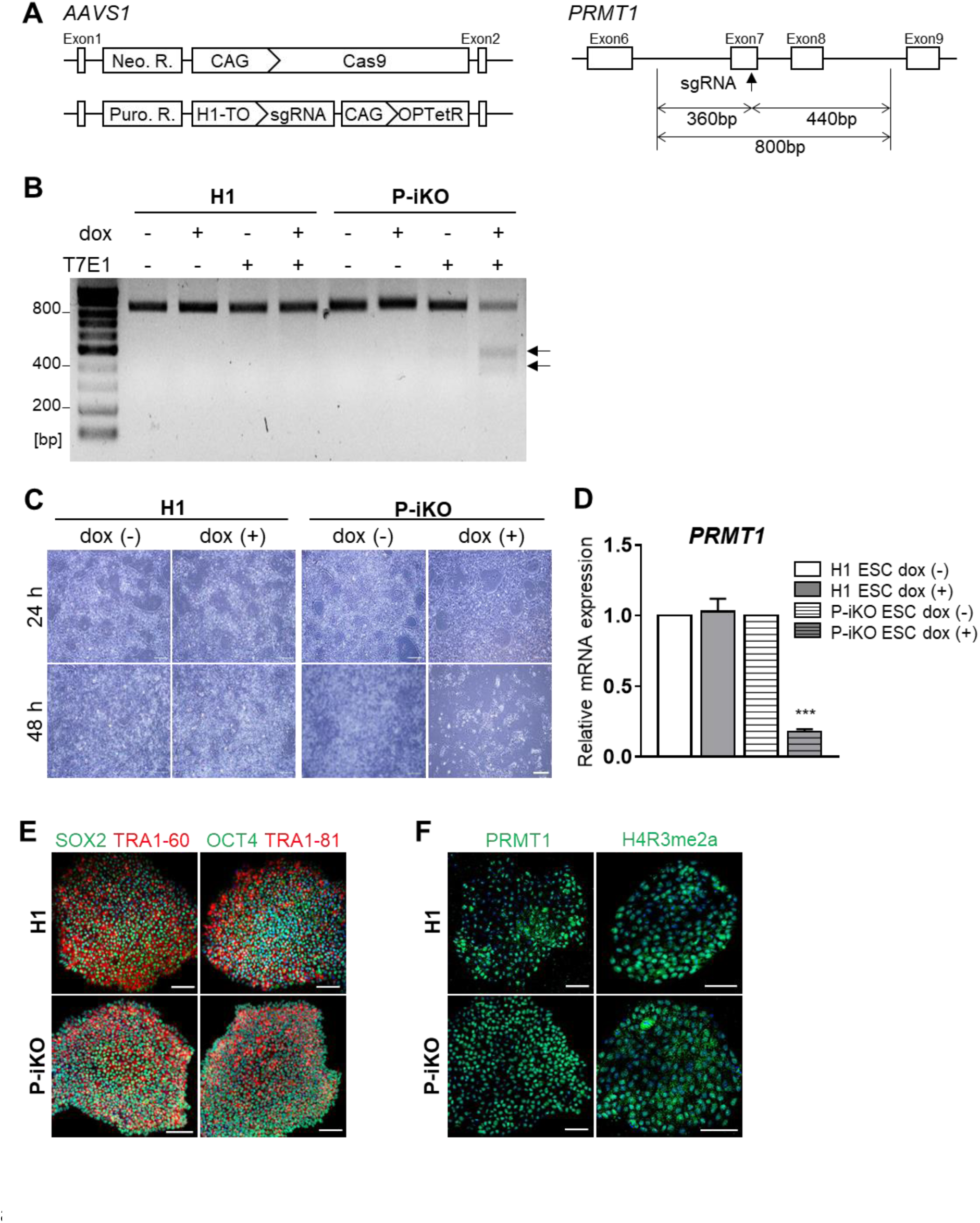
Generation of an P-iKO hESC line. (**A**) The *AAVS1* locus-targeted alleles for the generation of the P-iKO hESC line (left). Schematic of the *PRMT1* exon targeted using the doxycycline-inducible CRISPR system (right). See also Figure S1. (**B**) T7E1 analysis of genomic DNA from P-iKO hESCs. A single transgenic cell line after dox treatment was identified by the two genomic DNA fragments indicated by arrows (at 360 and 440 bp). (**C**) P-iKO hESC morphology. Most P-iKO hESCs died within 48 h of dox treatment. Scale bars, 500 μm. (**D**) Relative *PRMT1* mRNA levels in P-iKO hESCs. Relative expression values are presented as mean ± SEM (n = 3). \*\*\**p* < 0.001. (**E**) Normal expression of pluripotency-associated markers in P-iKO hESCs. Scale bars, 100 μm. (**F**) Immunostaining of PRMT1 and H4R3me2a in P-iKO hESCs. Scale bars, 100 μm.

### P-iKO hESCs show normal differentiation into pancreatic endocrine cells

The overall protocol we used to differentiate human ESCs into pancreatic endocrine cells (ECs) is depicted in Figure 2A. We found H1 and P-iKO hESCs both differentiated normally into definitive endoderm (DE), pancreatic endoderm (PE), endocrine progenitor (EP), and ECs in appearance (Figure S2A). In the absence of doxycycline (dox), we did not observe any difference between H1 and P-iKO hESCs in their transcription of genes associated with each developmental stage (Figure 2B). We also observed similar expression of the same markers in each developmental stage in an immunohistochemistry analysis comparing H1 and P-iKO hESCs (Figure 2C). In the absence of dox, the overall expression of PRMT1 was similar in each group, with PRMT1 expression falling significantly from the EP stage (Figures 2D and 2E). To determine the timing of dox treatment for P-KO induction, we examined *PDX1* transcriptional activity daily. In both H1 and P-iKO hESCs, *PDX1* transcripts gradually increased from day 3 of PE differentiation (PED3) and then decreased again upon reaching the EP stage (Figure 2F). During the same period, the transcriptional profiles of *PRMT1* also remained similar in both groups (Figure 2G). Thus, after confirming the similarity of H1 and P-iKO hESCs in the absence of dox, for all subsequent experiments, we added dox to the PE medium beginning on PED3 for an additional 3 d to knockout PRMT1 (Figure S2B). Together, these results confirm P-iKO hESCs show normal pluripotency and normal differentiation into pancreatic endocrine cells.

**Figure 2.**
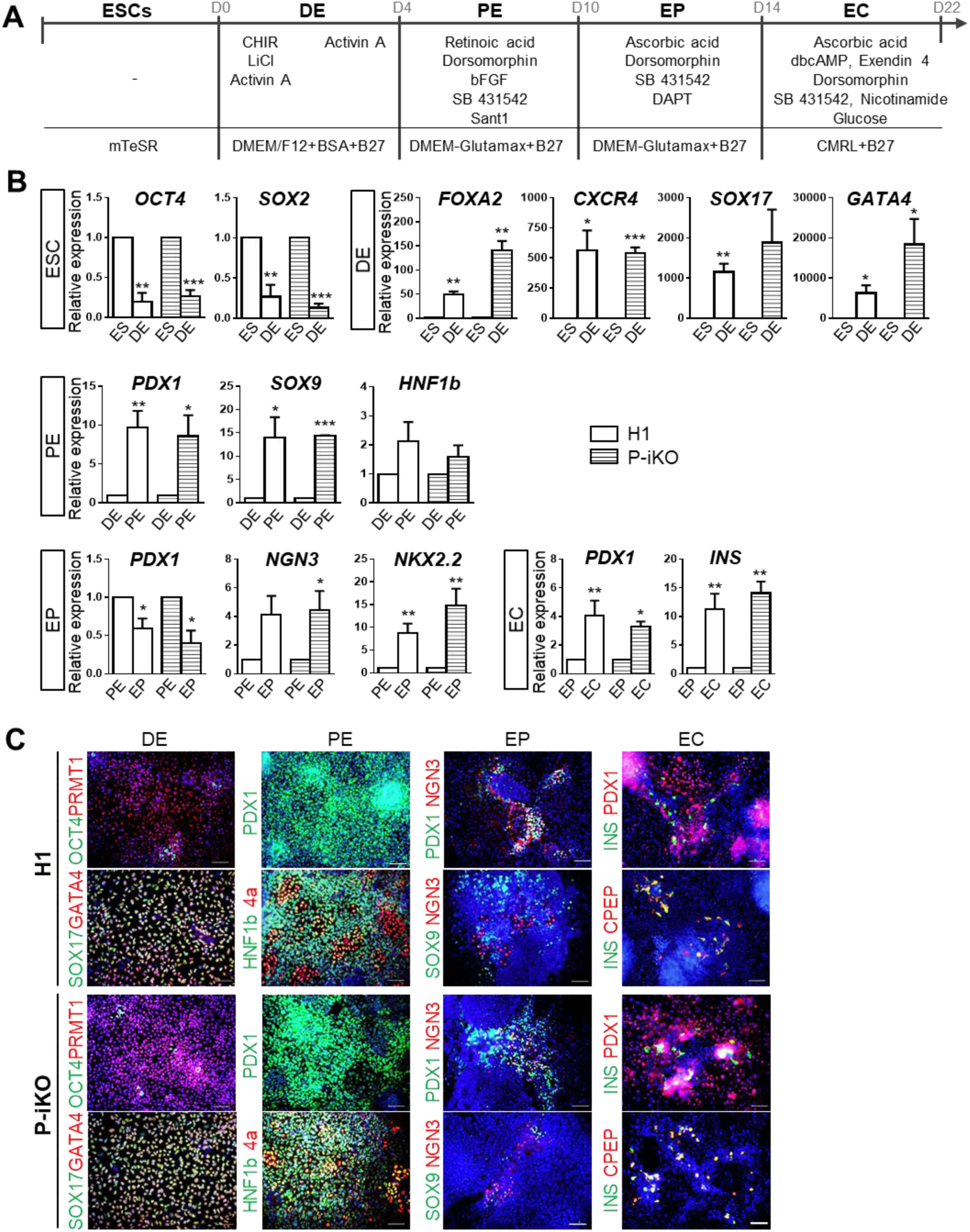
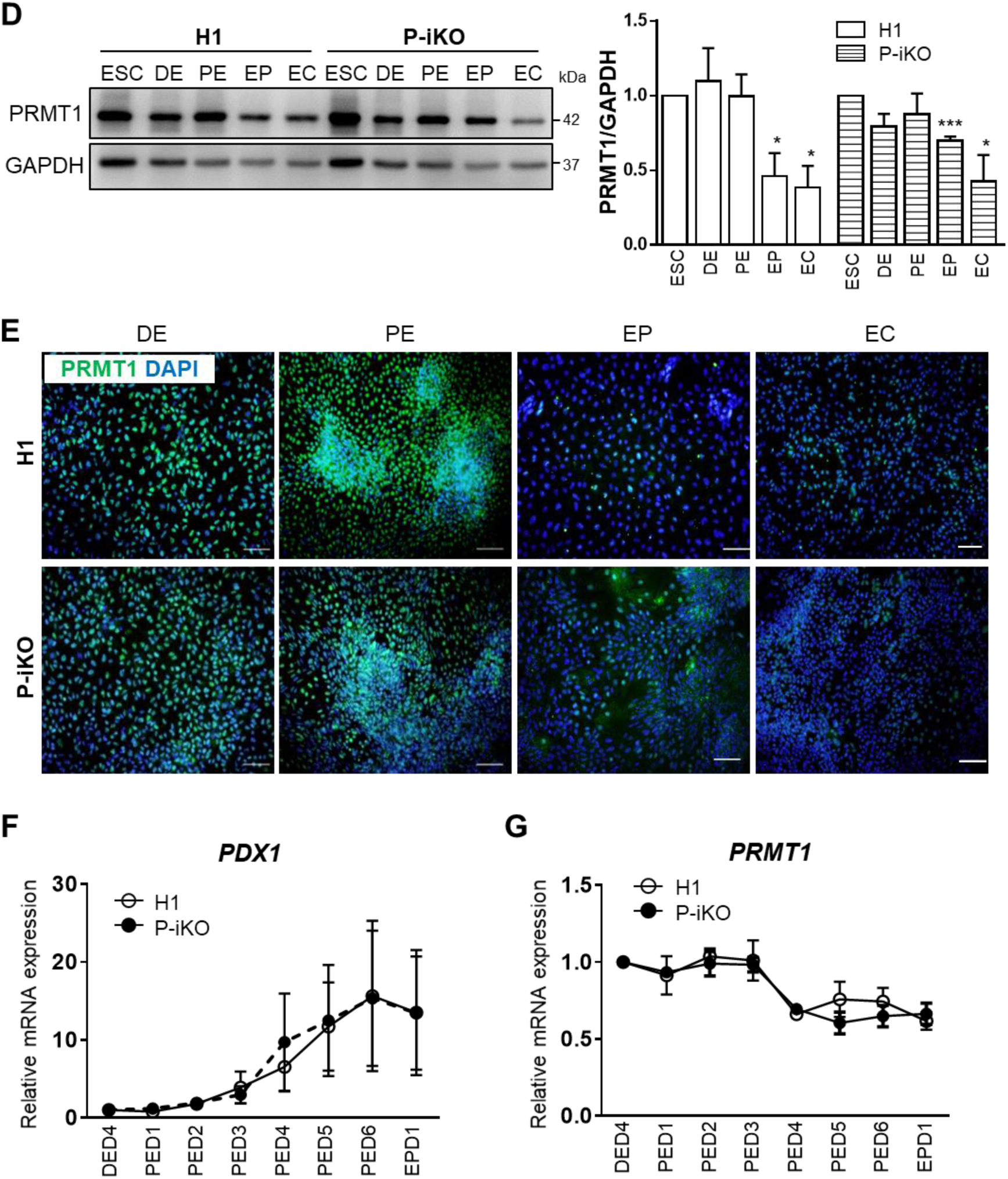
Differentiation of P-iKO hESCs into pancreatic endocrine cells. (**A**) Overall protocol for the differentiation of hESCs into pancreatic endocrine cells. Detailed procedures are described in the Materials and Methods section. D, day. See also Figure S2A. (**B**) Transcriptional expression of developmental genes in P-iKO hESCs in the absence of dox treatment during pancreatic differentiation, respectively. Relative expression values are presented as mean ± SEM. \**p* < 0.05, \*\**p* < 0.01, \*\*\**p* < 0.001 (n = 3–4). (**C**) Immunostaining of pancreatic developmental markers in P- iKO hESCs undergoing pancreatic differentiation. Scale bars, 100 μm. (**D**) Expression of PRMT1 protein in P-iKO hESCs as they progress through the stages of pancreatic differentiation. Relative expression values are presented as mean ± SEM. \**p* < 0.05, \*\*\**p* < 0.001 (n = 3). (**E**) Immunostaining of PRMT1 in P-iKO hESCs as they progress through the stages of pancreatic lineage differentiation. Scale bars, 100 μm. (**F**) Transcriptional profiles of the *PDX1* gene as P-iKO hESCs and controls progress from day 4 of the DE stage (DED4) to day 1 of the EP stage (EPD1) of pancreatic lineage differentiation. Data are represented as mean ± SEM (n = 6). (**G**) Transcriptional profiles of the *PRMT1* gene as P-iKO hESCs and controls progress from day 4 of the DE stage (DED4) to day 1 of the EP stage (EPD1) of pancreatic lineage differentiation. Data are represented as mean ± SEM (n = 5). See also Figure S2B.

### PRMT1-KO impairs the function of NGN3 in pancreatic endocrine development

NGN3 is an essential transcription factor for endocrine cell fate decision that has a very short half-life (Roark et al., 2012). We confirmed the short half-life of NGN3 in both H1 and P- iKO hESCs during *in vitro* pancreatic endocrine differentiation, observing robust expression in the EP stage that fell abruptly by the EC stage (Figure S2C). Another previous study showed accumulation of NGN3 in the pancreas of P-KO mice from E14.5 to the postnatal stage (Lee et al., 2019). To determine whether P-KO is also associated with NGN3 accumulation in this study, we treated P-iKO cells with dox in the PE stage for 3 d and further differentiated them into EP cells (Figure 3A). In the presence of dox, we observed reduced expression of PRMT1 (*p* < 0.001) but enhanced expression of NGN3 (*p* < 0.05) on day 4 of the EP stage (EPD4) of pancreatic endocrine differentiation (Figure 3B). We observed reduced asymmetrically di-methylated Histone 4 (H4R3me2a) in P-KO cells compared to untreated P-iKO cells (Figure 3B) and more NGN3-positive P-KO cells than untreated P-iKO cells (*p* <0.001, Figure 3C). This increase in NGN3-positive P-KO cells continued into the EC stage (Figure S3). Thus, we demonstrated that P-KO increases the proportion of NGN3-positive cells in human ESCs undergoing pancreatic development.

**Figure 3.**
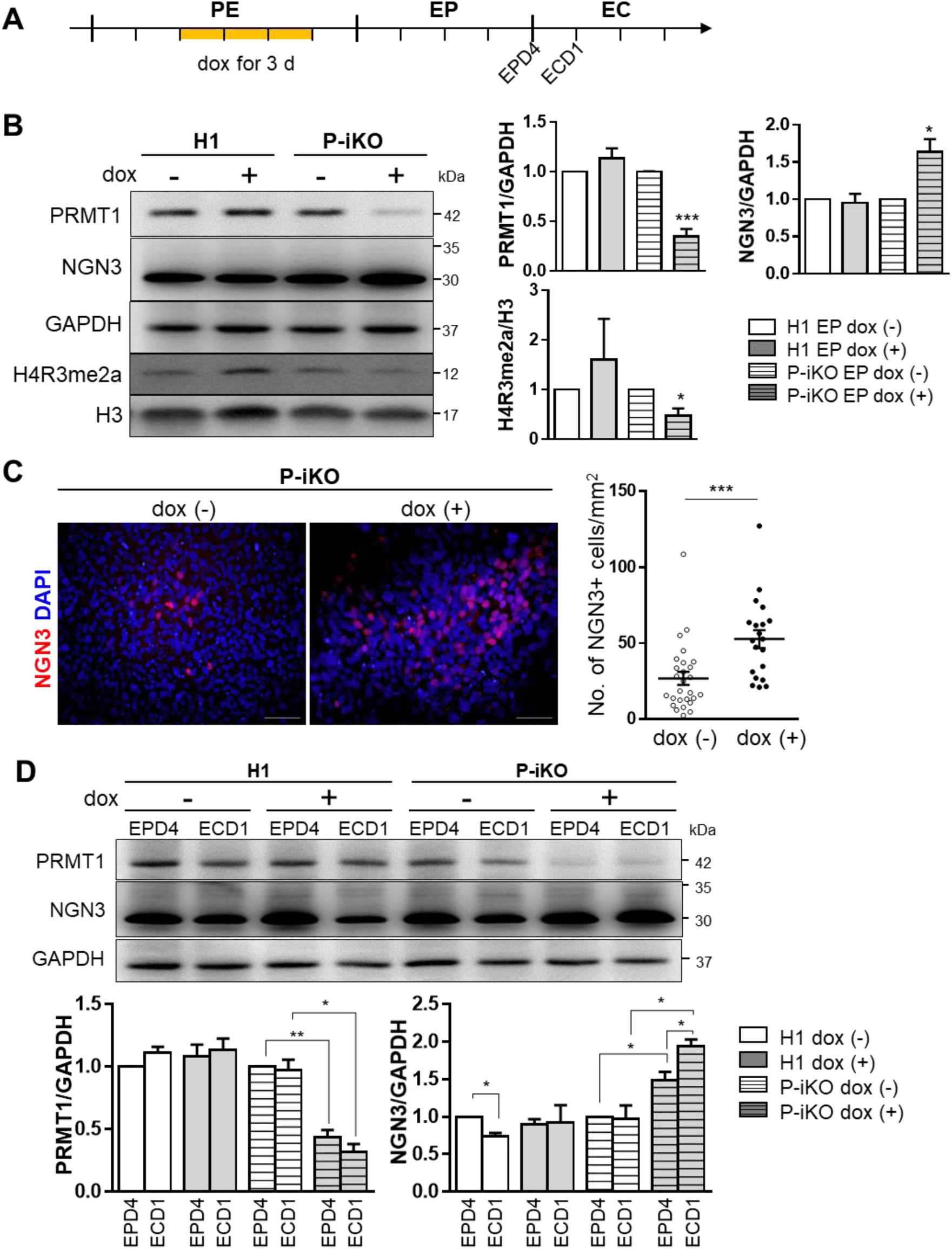
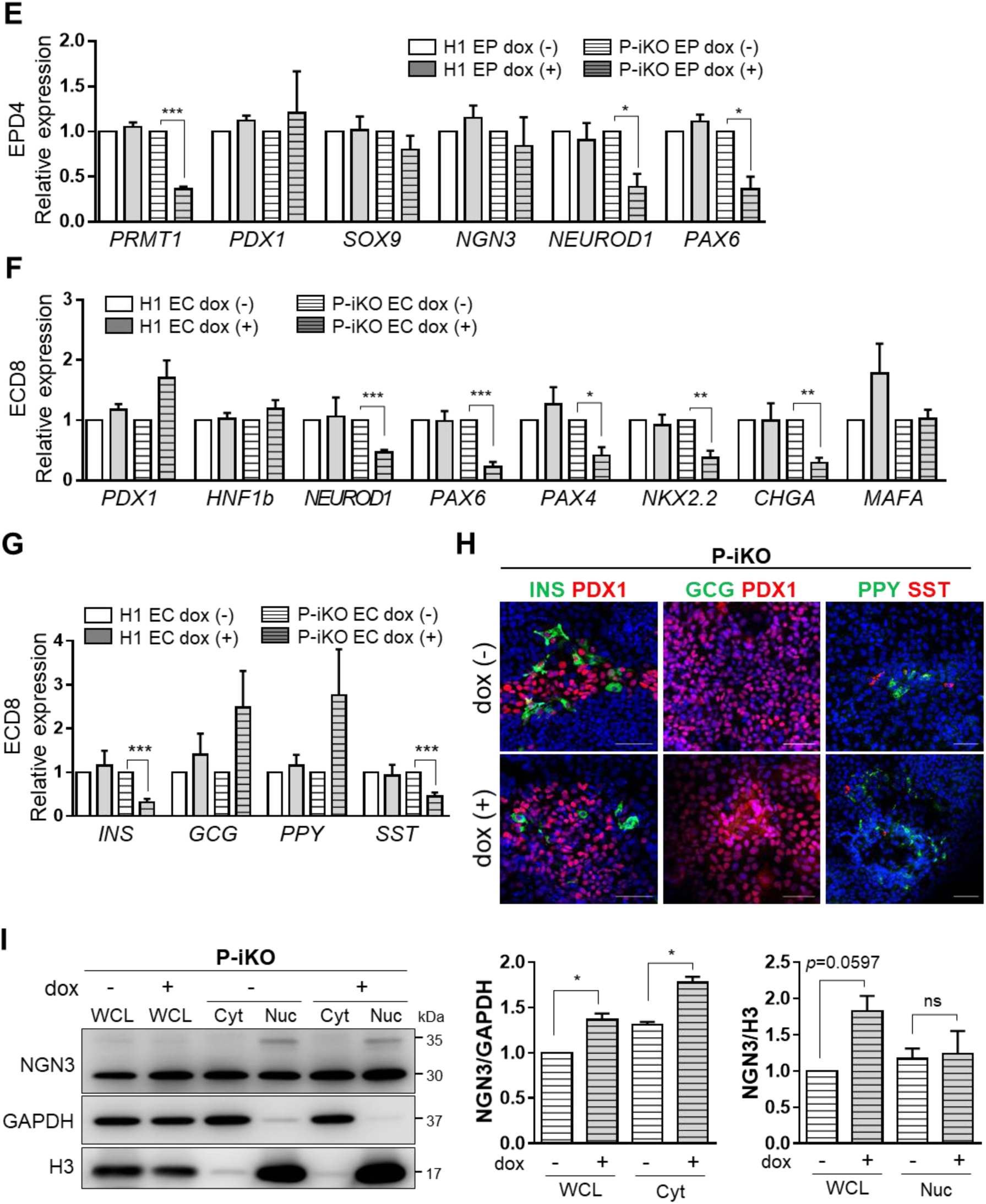
Effects of PRMT1-KO on the differentiation of pancreatic EPs to ECs. (**A**) Schedule for the dox treatment of PEs differentiated from P-iKO hESCs (yellow line). Dox was administered for 3 d to PRMT1-iKO hESCs that had progressed to D3 of the PE stage of pancreatic lineage development. (**B**) Expression of PRMT1 and NGN3 in P-iKO EP cells treated with dox. Relative PRMT1 and NGN3 protein expression levels were normalized to GAPDH levels. The H4R3me2a expression levels were normalized to H3 levels. Data are represented as mean ± SEM. \**p* < 0.05, \*\*\**p* < 0.001 (n = 3). (**C**) Immunostaining of NGN3 in P-iKO EP cells treated with dox (left). NGN3-positive cells were counted using ImageJ (right). P-KO EP cells accumulate NGN3. Data are represented as mean ± SEM. \*\*\**p* < 0.001 {dox (-), n = 27; dox (+), n = 20}. Scale bars, 50 μm. See also Figures S3A and S3B. (**D**) Expression of PRMT1 and NGN3 in P-iKO EP cells treated with dox. A western blot analysis showed reduced PRMT1 and enhanced NGN3 in P-KO EPD4 and ECD1 cells. PRMT1 and NGN3 protein expression levels were normalized by GAPDH. Dox (+) and (-) indicate treated and untreated cells, respectively. Data are represented as mean ± SEM. \**p* < 0.05, \*\**p* < 0.01 (n = 4). (**E**) Relative mRNA expression of EP- associated genes in P-KO EP cells. Transcription of NGN3 target genes was significantly downregulated in P-KO EP cells. Data are represented as mean ± SEM. \**p* < 0.05, \*\*\**p* < 0.001 (n = 4). (**F**) Relative mRNA expression of EC-associated genes in P-KO ECs. Transcripts of NGN3 target genes and EC-associated genes were significantly decreased in P-KO ECs. Data are represented as mean ± SEM.\**p* < 0.05, \*\**p* < 0.01, \*\*\**p* < 0.001 (n = 3). See also Figures S3C and S3D. (**G**) Relative mRNA expression of endocrine hormone genes in P-KO ECs. *INS* and *SST* genes were transcriptionally downregulated in P-KO ECs. Data are represented as mean ± SEM.\*\*\**p* < 0.001 (n = 4). INS, insulin; GCG, glucagon; PPY, pancreatic polypeptide; SST, somatostatin. (**H**) Immunostaining of endocrine hormones in P-KO ECs. Scale bars, 50 μm. (**I**) Cytoplasmic and nuclear fractionation of endogenous NGN3 in P-KO EP cells. P-KO EP cells accumulated NGN3 in the cytoplasm. Data are represented as mean ± SEM. WCL, whole cell lysate; Cyt, cytoplasm; Nuc, nucleus. \**p* < 0.05 (n = 3).

In general, NGN3 proteins rise in the EP stage and then fall in the EC stage as hESCs progress through pancreatic lineage differentiation (Figure S2C). Next, we asked whether P- KO is associated with NGN3 stability in the transition from the EP stage to the EC stage. Because we observed reduced PRMT1 expression in the EP and EC stages of P-iKO cells treated with dox compared to untreated cells, the same stages in which we saw increased NGN3 levels (Figure 3D), we hypothesized PRMT1 may play an important role in NGN3 stability. As a transcription factor, NGN3 activates the expression of pancreatic developmental genes, including *NEUROD1* and *PAX6* (Huang et al., 2000; Sander et al., 1997; Smith et al., 2004). We were surprised to find P-iKO cells treated with dox showed lower expression of *NEUROD1* and *PAX6* than untreated cells, although we did not detect any difference in the transcriptional activity of the pancreatic developmental genes *PDX1, SOX9*, or *NGN3* (Figure 3E). Considering the increased levels of NGN3 coupled with the reduced levels of NGN3 target genes (Figure 3D), it seems NGN3 does not activate its target genes in P-KO EP cells. We expected that either P-KO itself or a lack of arginine methylation is suppressing the functionality of NGN3 in hPSCs during pancreatic endocrine cell development.

Next, we investigated the developmental competence of P-KO EP cells to become pancreatic ECs. We found some EC-associated genes (i.e., *NKX2.2* and *CHGA*) with reduced expression in P-KO ECs and others (i.e., *PDX1, HNF1b*, and *MAFA*) with normal expression (Figure 3F). Transcripts of genes downstream of NGN3 (i.e., *NEUROD1, PAX6*, and *PAX4*) were reduced in P-KO ECs (Figure 3F). Among the pancreatic endocrine hormone genes, we found specific reductions in insulin (INS) and somatostatin (SST) expression at the mRNA and protein levels in P-KO ECs (Figures 3G and 3H). Our results imply PRMT1 KO leads to reduced transcriptional activity of NGN3, thereby impairing the competence of EP cells to become β- or δ-cells *in vitro*. To determine whether P-KO influences the cellular localization of NGN3, we examined NGN3 expression in the cytoplasmic and nuclear compartments of P-iKO EP cells. When compared to untreated cells, dox-treated P-iKO EP cells showed higher levels of NGN3 protein in whole cell lysates (WCL) and in the cytoplasm (Cyt) but no difference in the nuclear compartment (Figure 3I). On western blot, we observed multiple nuclear NGN3 bands in P-iKO EP cells but only a single cytoplasmic NGN3 band. Thus, NGN3 accumulates in the cytoplasm of P-KO EP cells. These data suggest PRMT1 KO causes NGN3 accumulation in EP cells, leading to impaired EC differentiation.

### PRMT1 specifically methylates NGN3 arginine 65

PRMT1 preferentially methylates the arginine residues of RGG/RXR sequences in glycine and arginine-rich (GAR) motifs in histone and non-histone proteins (Bedford and Richard, 2005; Gary and Clarke, 1998; Thandapani et al., 2013). By checking UniProt (www.uniprot.org), we found NGN3 has a highly conserved RGG motif (amino acids 65–67) near its basic helix-loop-helix (bHLH) domain (aa 83–135) (Figure 4A). To determine whether arginine methylation of NGN3 is regulated by PRMT1, we performed a co-immunoprecipitation (co-IP) assay to investigate physical contact between PRMT1 and NGN3. First, we made various FLAG- or GST-tagged NGN3 expression vectors containing fragments of the NGN3 gene. These included vectors containing intact NGN3, NGN3 fragment 1 (aa 1–80, f1), NGN3 fragment 2 (aa 81–214, f2), and an NGN3 R65A mutant (Figure 4B). We generated this NGN3 R65A mutant with an alanine (GCC) replacing arginine 65 (CGC) in hopes that PRMT1 would not be able to methylate it. To verify the interaction of NGN3 with PRMT1, we transfected pCAG-FLAG-NGN3 WT or pCAG-FLAG-NGN3 R65A mutant plasmids into HEK cells and then performed an immunoprecipitation experiment. The transfectants expressed endogenous PRMT1 and either FLAG-NGN3 WT or R65A mutant (Figure 4C, left two lanes). We were only able to detect mono- and di-arginine methylation bands (arrows) in the FLAG-NGN3 WT eluate (Figures 4C and 4D). This result indicates PRMT1 methylates NGN3 arginine 65. Thus, we decided to focus on arginine 65 methylation to clarify its role in trans-activation.

**Figure 4.**
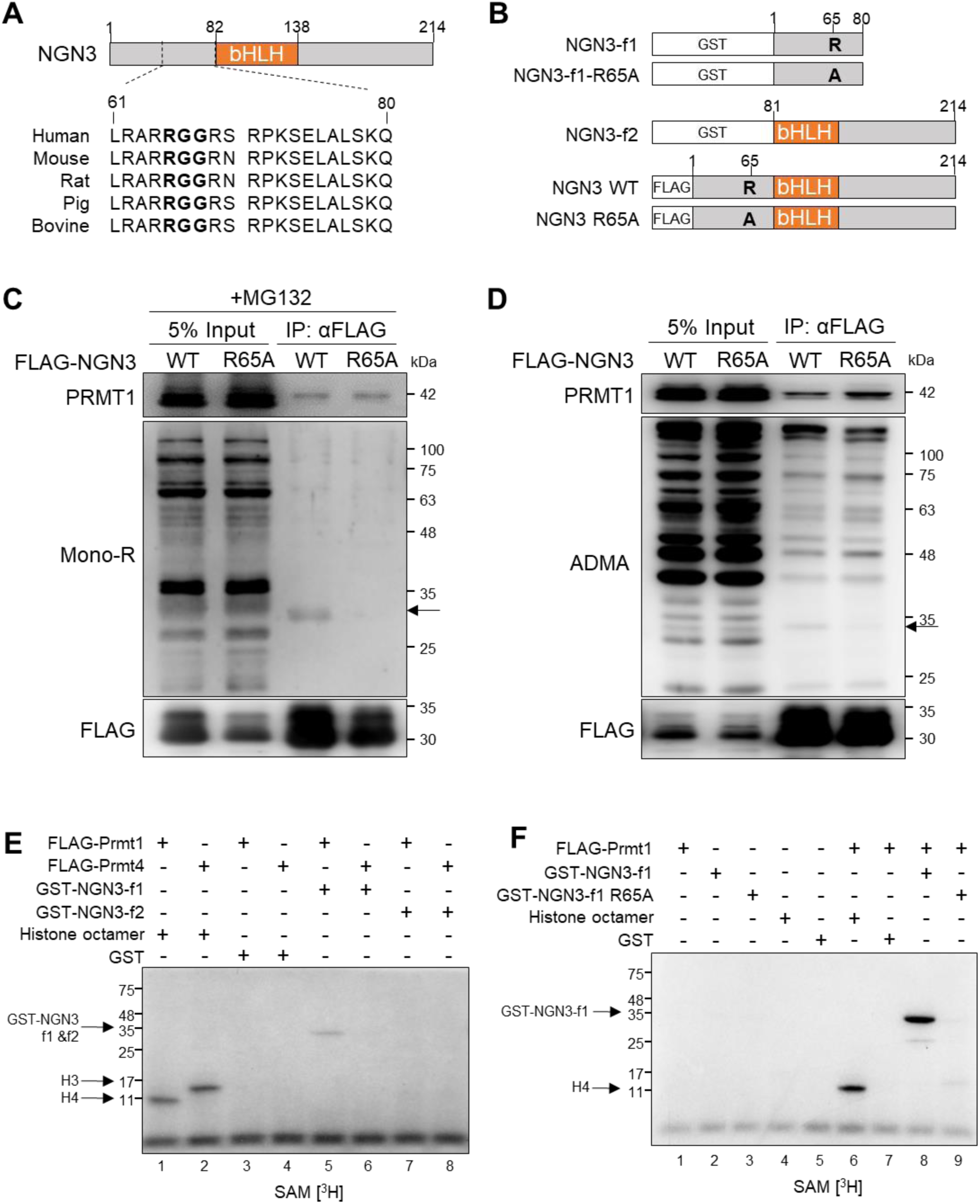
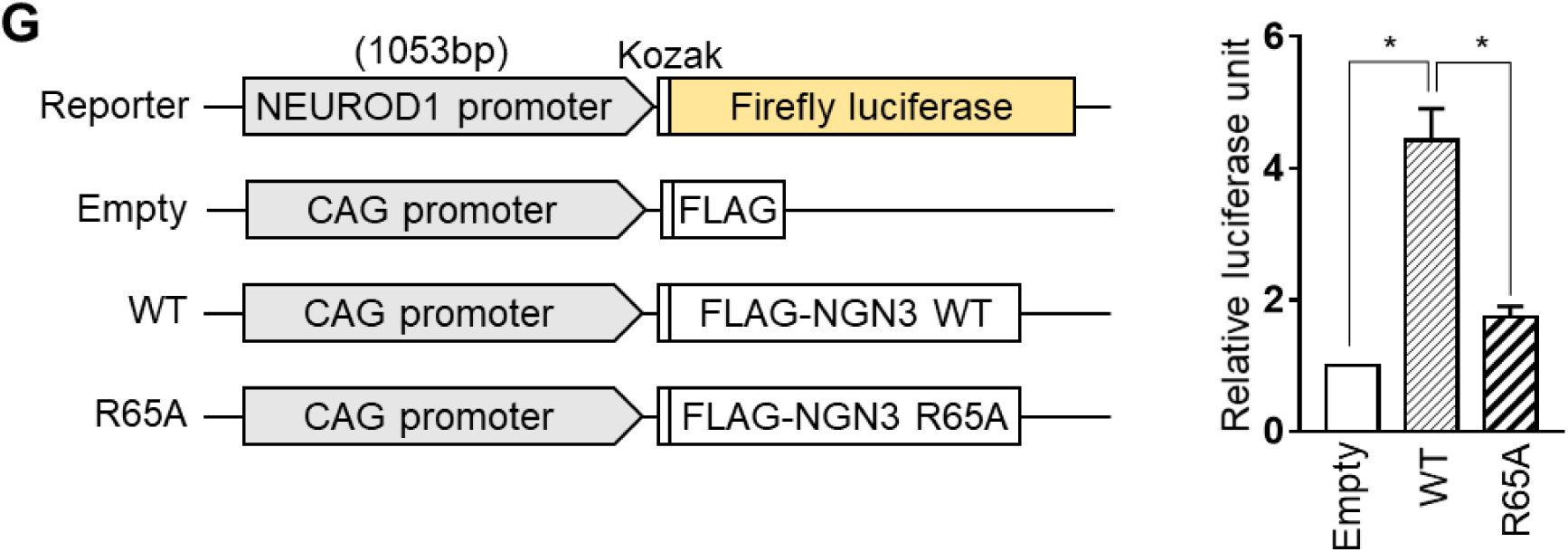
PRMT1-mediated methylation of NGN3 arginine 65. (**A**) An RGG motif (bold) in human NGN3 is conserved across species. (**B**) Several human NGN3 constructs for arginine methylation analysis. GST-tagged NGN3 fragments 1 (f1) and 2 (f2) were inserted into pGEX4T-1 for methyl transferase assays. FLAG-tagged NGN3 constructs were inserted into pCAG-FLAG-IPuro for NGN3 overexpression. Arginine 65 [R] of NGN3 was changed to alanine [A] to prevent arginine methylation. (**C**) Immunoprecipitation of FLAG-NGN3 with MG132 treatment. A mono-R band (arrow) was detected for FLAG-NGN3 WT but not for FLAG-NGN3 mutant. mono-R, mono-methylated arginine. (**D**) Immunoprecipitation of FLAG-NGN3 without MG132 treatment. A di-methylated ADMA band was only detected with FLAG-NGN3 WT. ADMA, asymmetrical di-methylated arginine. (**E**-**F**) In vitro NGN3 methylation assays. NGN3 arginine 65 was methylated in the presence of [^3^H] SAM by Prmt1 (E, lane 5). The R65A NGN3 mutant was not methylated in the presence of [3H] SAM by Prmt1 (F, lane 8). Histone octamers were used to test FLAG-PRMT1 and FLAG-Prmt4 activities. The GST tag served as a negative control. See also Figure S4. (**G**) Luciferase assay for NGN3 activity in HEK cells using the *NEUROD1* promoter. The firefly luciferase signal for each experiment was normalized to the renilla luciferase signal. Data are presented as mean ± SEM. \**p* < 0.05 (n = 3). See also Figure S5.

First, we purified FLAG-Prmt family members and a series of recombinant GST-NGN3 fragments for use in methyl transferase assays (Figure S4A). While both FLAG-Prmt1 and FLAG-Prmt4 could methylate histone octamers, only Prmt1 and not Prmt4 could methylate GST-NGN3-f1 (Figure 4E). We were also unable to observe any methylation signal at all with GST-NGN3-f2 (Figure 4E, lanes 7 and 8). In contrast to our results with WT-NGN3 (Figure 4F, lane 8), the R65A-NGN3 fragment 1 seemed to prevent methylation (Figure 4F, lane 9). Both mono- and dimethylation of arginine showed Prmt1 dose-dependence (Figure S4C). Of all the Prmt family members, we only observed significant methylation of the WT-NGN3 fragment with Prmt1 (Figure S4B, arrow). Even this did not occur, however, in the combination of PRMT1 and R65A-NGN3-f1. Thus, our results indicate PRMT1 specifically methylates NGN3 arginine 65 *in vitro*.

Next, we asked whether arginine methylation of NGN3 influences the expression of its downstream genes. After transfecting FLAG-tagged WT-NGN3 and R65A-NGN3 into HEK cells, we found R65A-NGN3 triggered less activation of *NEUROD1* than WT-NGN3 (Figure S5A). We then confirmed this reduced transcriptional activity of the R65A-NGN3 mutant via luciferase assay. To do so, we co-transfected HEK cells with each expression vector and a firefly luciferase reporter containing the *NEUROD1* promoter (Huang et al., 2000; Wang et al., 2006). We first confirmed that the expression of each FLAG-tagged version of NGN3 was similar (Figure S5B) and then normalized the firefly luciferase signal to the renilla luciferase signal. As expected, we observed significantly higher activity triggered by the WT-NGN3 transfectants than the R65A-NGN3 transfectants (Figure 4G, p<0.05). This result indicates that the R65A mutation blocks NGN3 transcriptional activity. Together, our findings suggest methylation of NGN3 arginine 65 is essential for its transcriptional activity.

### Arginine 65 methylation of NGN3 is a prerequisite for its degradation

NGN3 is transiently expressed in pancreatic EP cells, rapidly disappearing during pancreatic lineage development (Apelqvist et al., 1999). We therefore asked whether arginine methylation of NGN3 is associated with its degradation. To address this question, we transfected P-iKO PE cells with an NGN3 overexpression vector (pCAG-FLAG-NGN3) and incubated the transfected cells for 1–2 d (Figure 5A). Although ectopic NGN3 was significantly reduced 2 days after transfection in P-iKO PE cells that were not treated with doxycycline, similar cells treated with doxycycline maintained consistent levels of NGN3 (Figure 5B). This suggests PRMT1 is associated with NGN3 degradation in hESCs during pancreatic lineage differentiation.

**Figure 5.**
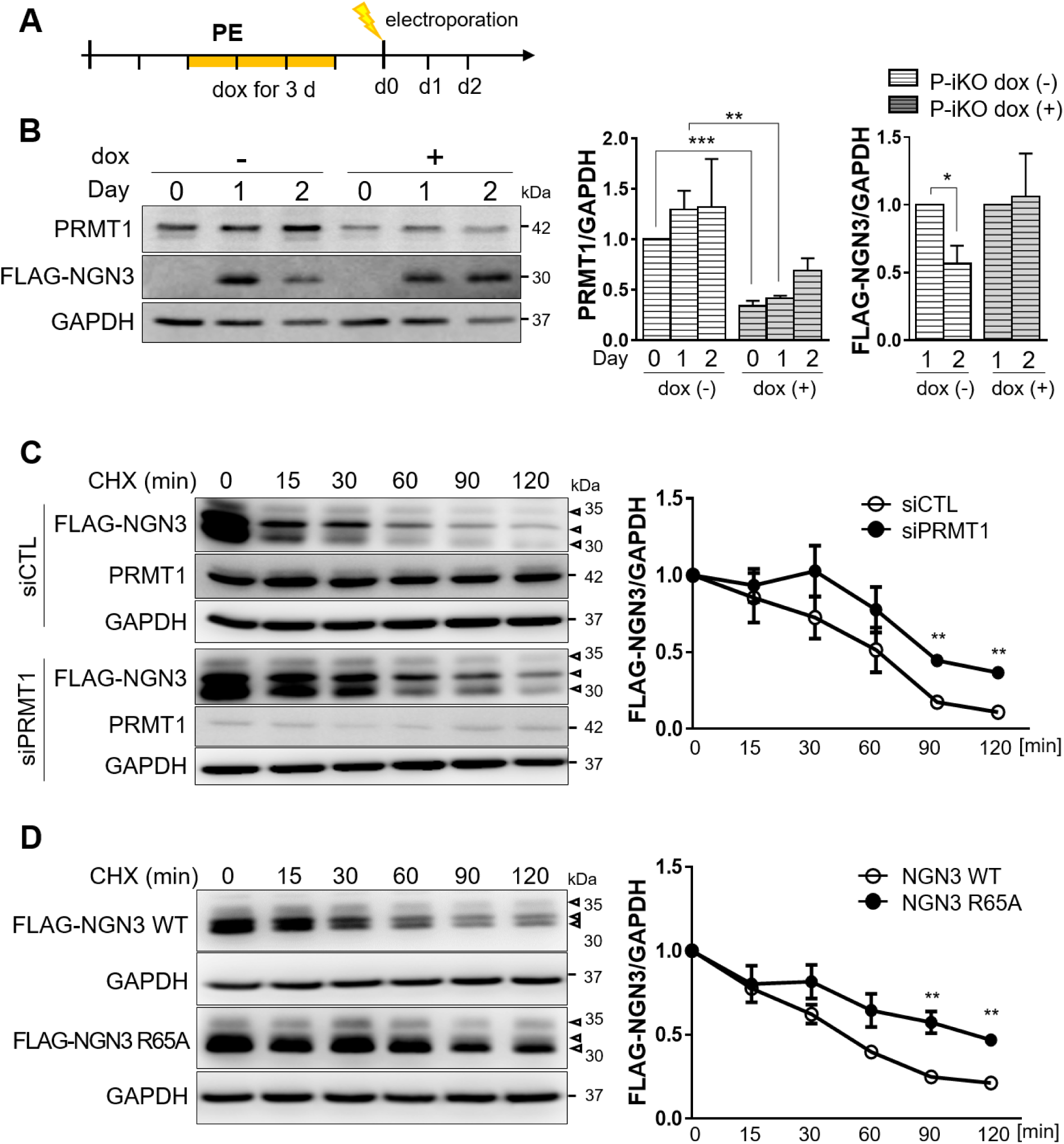
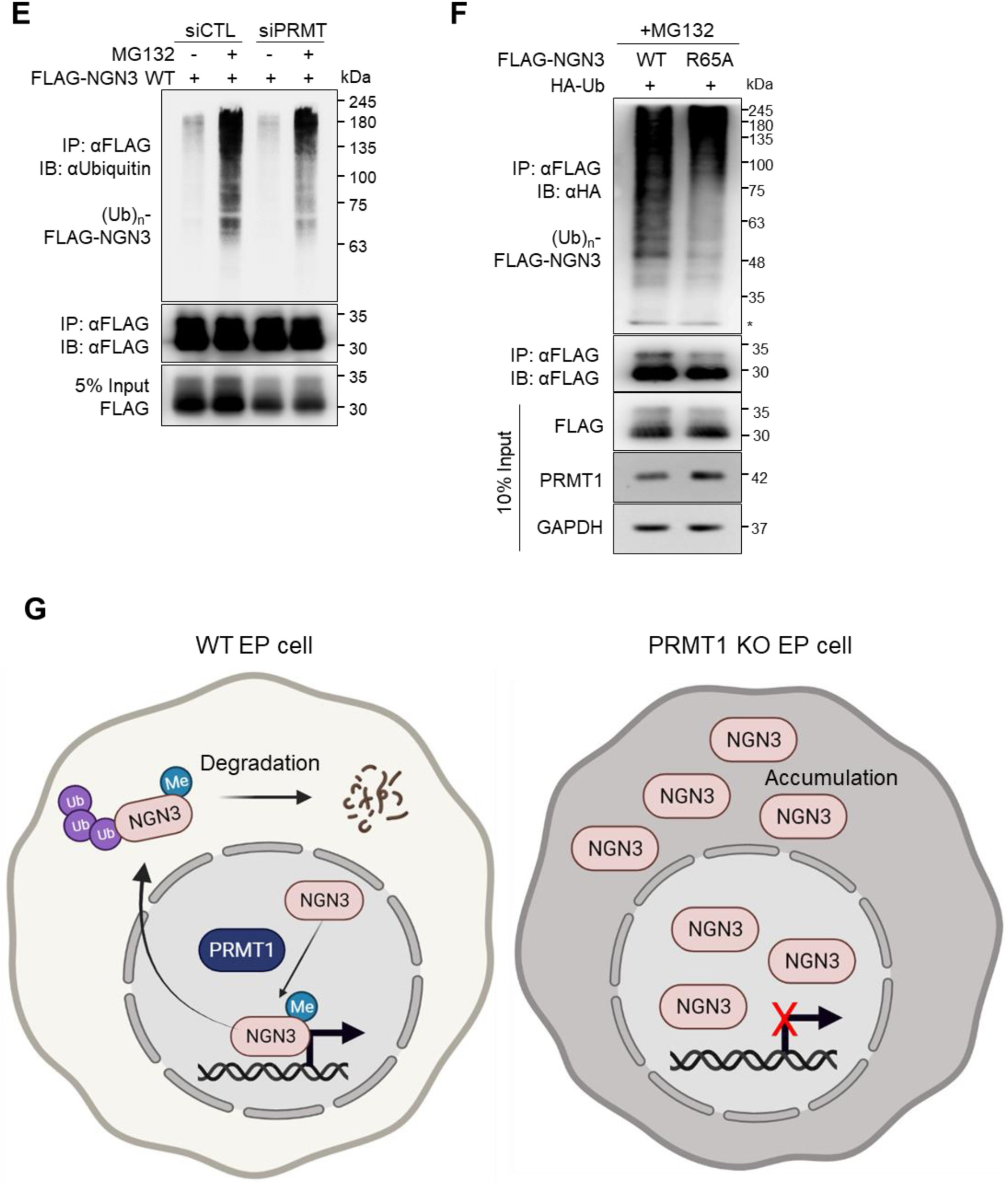
Degradation of arginine 65-methylated NGN3. (A) Protocol for the treatment of P-iKO PE cells with dox and electroporation with the FLAG-NGN3 expression vector. (B) Accumulation of NGN3 protein in P-KO PE cells. P-iKO PE cells treated with dox (+) showed a downregulation of PRMT1 but not FLAG-NGN3. Data are presented as mean ± SEM. *p < 0.05, **p < 0.01, ***p < 0.001 (n = 4). (C) Degradation assay for FLAG-NGN3 in HEK cells treated with PRMT1-siRNA. In the presence of CHX, we observed delayed degradation of FLAG-NGN3 in PRMT1-KD HEK cells. FLAG-NGN3 proteins appear as multiple bands (arrowheads). FLAG- NGN3 levels in cells treated with siPRMT1 compared to those treated with siCTL at the indicated time points after CHX treatment. Data are presented as mean ± SEM. **p < 0.01 (n = 3). CHX, cycloheximide; siCTL, siRNA control (unspecific sequences). (D) Delayed degradation of FLAG-NGN3 R65A mutant in HEK cells. FLAG-NGN3 R65A mutant was degraded slower than FLAG-NGN3 WT. FLAG-NGN3 R65A mutant levels compared to FLAG-NGN3 WT at the indicated time points after CHX treatment. Data are presented as mean ± SEM. **p < 0.01 (n = 3). See also Figure S6. (E) Reduced polyubiquitination of FLAG-NGN3 in PRMT1-KD HEK cells. HEK cells transfected with PRMT1-siRNA showed reduced ubiquitination. (F) Decreased polyubiquitination of FLAG-NGN3 R65A mutant in HEK cells. (G) Putative model for the role of arginine 65 methylation of NGN3 in pancreatic EP cells. Arginine- methylated NGN3 activates NGN3 target genes like NEUROD1 in the nucleus and is then rapidly degraded by ubiquitination at the pancreatic EP stage. PRMT1-KO prevents NGN3 arginine methylation, leading to the accumulation of NGN3 and reduced activation of its target genes. Ultimately, defective NGN3 arginine 65 methylation prevents the cell fate transition of EPs to ECs. The diagram was generated using Biorender (biorender.com).

To investigate the role PRMT1 plays in NGN3 degradation, we transfected HEK cells with PRMT1-specific siRNAs, waited 48 h, further transfected them with a FLAG-NGN3 expression vector, waited another 24 h, and then treated them with cycloheximide (CHX) to block protein synthesis. We observed a significant delay in FLAG-NGN3 degradation in the siPRMT1 group compared to the siCTL group (Figure 5C, *p* < 0.001). We also found that FLAG-NGN3 R65A showed a longer half-life than FLAG-NGN3 WT (Figure 5D, *p* < 0.001). Unlike R65A-NGN3, WT-NGN3 appeared in multiple distinct bands on western blot (Figure 5D). In addition, we observed differences in the various phosphorylation bands for FLAG-tagged WT-NGN3 and R65A-NGN3 (Figure S6A, arrowheads) that were sensitive to phosphatase treatment (Figure S6B, arrowheads). These results indicate arginine methylation contributes to NGN3 stability by altering its phosphorylation.

To examine the correlation between arginine methylation and polyubiquitination of NGN3, we transfected HEK cells with FLAG-tagged WT-NGN3 plasmids and siPRMT1 and then treated the transfected cells with the proteasome inhibitor MG132 (20 μM) for 2 h. In the presence of proteasome inhibitor, the ubiquitinated NGN3 bands were slightly reduced in the siPRMT1 group compared to the siCTL group (Figure 5E). R65A-NGN3 also showed reduced ubiquitination bands compared to FLAG-tagged WT-NGN3 (Figure 5F). These results suggest PRMT1-mediated arginine methylation is associated with NGN3 polyubiquitination. Together, our results indicate PRMT1 specifically methylates arginine 65 of NGN3 in hESCs during pancreatic EC development. This methylated NGN3 then undergoes ubiquitin-mediated degradation.

## Discussion

Here, we report for the first time that arginine 65 methylation of NGN3 is essential in the pancreatic lineage differentiation of hESC-derived EP cells to ECs. While we did find that PRMT1-KO (P-KO) EP cells exhibit increased NGN3 protein levels, they do not develop into pancreatic ECs *in vitro*. Using an in vitro methylation assay, we found PRMT1 specifically methylates arginine 65 on the GAR motif of NGN3. Preventing the methylation of arginine 65 blocked ubiquitin-mediated NGN3 degradation. Thus, we found PRMT1 is a key regulator of NGN3 function in pancreatic development via direct PRMT1-NGN3 crosstalk.

Arginine methylation of non-histone proteins is important in several cellular functions, such as transcription, EMT, and cellular proliferation (Avasarala et al., 2015; Bedford and Clarke, 2009; Gary and Clarke, 1998; Raposo and Piller, 2018; Yamagata et al., 2008). The role of PRMT1-mediated arginine methylation in pancreas development, however, is poorly understood. PRMT1-KO (PKO) mouse embryos exhibit prolonged NGN3 expression accompanied by reduced pancreatic β-cell mass and pancreatic hypoplasia after birth (Lee et al., 2019). This suggests PRMT1 may control NGN3 stability in mammalian pancreatic development. Transient expression or rapid degradation of NGN3 is necessary for cell fate determination of EPs into ECs (Gradwohl et al., 2000; Miyatsuka et al., 2011; Salisbury et al., 2014; Smith et al., 2004). To study the molecular mechanism underlying PRMT1’s role in NGN3 stability as it relates to pancreatic development, we generated a P-iKO hESC line and guided its differentiation into the pancreatic endocrine lineage. Despite producing increased levels of NGN3, P-KO EP cells did not develop into ECs (Figure 3). Furthermore, we found P-KO EPs and ECs exhibited reduced expression of various genes downstream of NGN3 (i.e., *NEUROD1, PAX6, PAX4, NKX2.2,* and *CHGA*). These results suggest P-KO EP cells have significant NGN3 dysfunction. PRMT1 deficiency affects cell fate decisions in specialized endocrine cell types, as P-KO ECs show significantly reduced expression of β- and δ-cell markers (Figure 3G). Since β- and δ- cells arise from the same ancestor (DiGruccio et al., 2016; Mfopou et al., 2010), it is clear that PRMT1-KO contributes to the specification of EP cells into β- and δ- cells. We found that arginine 65 of NGN3 is a conserved methylation motif among mammals whose methylation is specifically catalyzed by PRMT1 (Figure 4). Furthermore, unmethylated NGN3 was not degraded via ubiquitin-mediated proteolysis (Figures 5E and 5F). Our results indicate NGN3 arginine 65 methylation is crucial for EC fate determination in pancreatic lineage development.

To induce PRMT1 loss of function at a specific stage of pancreatic development, we generated a PRMT1-inducible KO (P-iKO) system in hESCs using a “safe harbor” *AAVS1* locus. We selected this *AAVS1* locus because it does not produce any adverse effects on differentiation, transgene silencing, or proliferation in gene-edited hiPSCs (Hockemeyer et al., 2011; Smith et al., 2008). We performed homology-directed repair (HDR) using ZFNs to avoid random transgene integration. When we cultured P-iKO hESCs in the presence of doxycycline, most died within 2 d (Figure 1C). This implies PRMT1 is essential for the viability and proliferation of hESCs, which is consistent with the embryonic lethal phenotype of PRMT1-KO mice (Pawlak et al., 2000). In the absence of dox, however, P-iKO hESCs maintained pluripotency, differentiation competence, and Cas9 mRNA expression for more than 3 months (data not shown). Thus, our use of the *AAVS1* locus with these transgenes permitted stable enough gene expression for the maintenance and differentiation of hESCs.

NGN3 protein is rapidly degraded by the canonical ubiquitination pathway (Vosper et al., 2009). In the cytoplasm of mouse EPs, NGN3 is degraded by the ubiquitin-proteasome system after phosphorylation (Krentz et al., 2017; Vosper et al., 2009). As in mouse EPs, we observed multiple NGN3 phosphorylation bands in the nuclear fraction of EP cells derived from hESCs (Figure 3I). We observed a different electrophoretic mobility for the NGN3 R65A mutant compared to WT NGN3 (Figure S6, arrowheads), suggesting arginine 65 methylation of NGN3 affects its phosphorylation. In addition, we found that most of the endogenous NGN3 protein in P-KO EP cells was localized to the cytoplasm (Figure 3I). As with the NGN3 R65A mutant (Figure 5D), KD of PRMT1 slowed NGN3 degradation in HEK cells (Figure 5C). Therefore, we concluded arginine 65 methylation of NGN3 is associated with NGN3 phosphorylation, leading to NGN3 degradation. This degradation is required for the cell fate transition of EPs to ECs during human pancreatic lineage development.

From these results, we propose a model for the role of PRMT1 in pancreatic endocrine lineage development (Figure 5G). In this model, NGN3 arginine 65 is specifically methylated by PRMT1. This permits the expression of genes downstream of NGN3 that are important for the transition of EPs to ECs. The methylation of NGN3 arginine 65 also induces ubiquitin-dependent degradation of NGN3 by the proteasome, permitting the completion of EC development. In the absence of PRMT1, NGN3 does not activate the expression of its downstream target genes, nor is it degraded to permit the EP-to-EC transition. Therefore, we conclude arginine 65 methylation of NGN3 is essential for EC fate determination during pancreatic development.

## Materials and Methods

### Plasmid constructs for the PRMT1-inducible Knockout (P-iKO) system and for NGN3 expression

The ZFN-L, ZFN-R, pAAV-Neo_Cas9, and pAAV-Puro_siKO vectors (Bertero et al., 2016; Snijders et al., 2019) were provided by Prof. KJ Yoon at the Department of Biological Sciences, KAIST. The pAAV-Puro_siKO vector was introduced a chimeric sgRNA expression cassette using the AarⅠ restriction enzyme (New England Biolabs, Ipswich, MA). The sgRNA 5’-ATGTGACGGCCATCGAGGAC-3’ and PAM CGG were selected from several sequences recommended by the IDT CRISPR_PREDESIGN program (Integrated DNA Technologies, Inc., IA). For the expression of human NGN3, either a full-length NGN3 cDNA or NGN3 cDNA fragments were inserted into the pCAG-FLAG-IPuro or pGEX4T-1 vectors, respectively. Site-directed mutagenesis (R65A) was performed by Bioneer (BIONEER Co., Daejeon, Korea). All plasmids were cloned into the DH5a strain (Enzynomics) and purified with a NucleoBond Xtra Midi Plus kit (MACHEREY-NAGEL GmbH & Co. KG, Dueren, Germany) according to the manufacturer’s instructions.

### Generation of the P-iKO human embryonic stem cell (hESC) line

H1 hESCs were transfected with ZFN-L (5 μg), ZFN-R (5 μg), pAAV-Neo_Cas9 (2.5 μg), and pAAV-Puro_siKO (2.5 μg) plasmids with a NEPA21 Super Electroporator (Nepa Gene, Chiba, Japan). A 150 V poring pulse and a 20 V transfer pulse were used for electroporation. Transfected cells were placed on a Matrigel-coated culture dish and incubated in mTeSR1 medium (STEMCELL Technologies Inc., Vancouver, Canada) containing 10 μM Y27632 (ROCK inhibitor, A. G. Scientific, San Diego, CA) at 37℃ and 5% CO_2_ for 3 d until they reached confluence. For transgenic colony selection, the cells were treated with 200 μg/ml G418 (InvivoGen, Pak Shek Kok, Hong Kong) for 4 d and then with 0.5 μg/ml puromycin (Sigma, St. Louis, MO) for 3 d. Finally, a single colony with an sgRNA cassette and a Cas9 transgene at the AAVS1 locus (P-iKO hESC line) was obtained for this study. Wild type (WT) and P-iKO hESCs were maintained in mTeSR1 medium on Matrigel-coated plates at 37℃ and 5% CO_2_ with daily media changes. To expand the hESCs, single ESC colonies were mechanically divided into 20–25 clumps. These clumps were treated with 10 μg/ml Dispase (Invitrogen, Carlsbad, CA) at 37℃ for 4 min, divided into 3–4 culture dishes, and incubated under the conditions described above. For use in subsequent and repetitive experiments, some transgenic cell clumps from the early passages were frozen in KnockOut serum (Invitrogen) containing 10% DMSO.

### Differentiation of hESCs into pancreatic endocrine cells

Human ESCs were differentiated into pancreatic endocrine cells as previously reported (Kim et al., 2016). During differentiation of the pancreatic endocrine lineage, ESCs pass through definitive endoderm (DE), pancreatic endoderm (PE), endocrine progenitor (EP), and endocrine cell (EC) stages. Briefly, H1 and P-iKO hESC colonies were dissociated into single cells with 0.5 μM EDTA at 37℃ for 6 min and then harvested. Approximately 6.5×10^4^ live cells were incubated on Matrigel-coated 4-well plates (SPL lifesciences, Pocheon, Korea) in 500 μl of mTeSR1 medium supplemented with 10 μM Y27632 at 37℃ and 5% CO_2_ for 2 d. To induce DE formation, the resulting cells were incubated in DMEM/F12 medium supplemented with 0.2% BSA (Sigma), 50 ng/ml activin A (R&D Systems, Inc., Minneapolis, MN), 3 μM CHIR99021 (Cayman Chemical, Ann Arbor, MI), and 2 mM LiCl (Sigma) for 1 d. Then, they were further cultured in DMEM/F12 supplemented with 0.2% BSA (Sigma), 1% B27 supplement (Invitrogen), and 50 ng/ml activin A (R&D Systems) for 3 d. DMEM/F12 medium is composed of 1.2 g/L sodium bicarbonate (Sigma), 1 mM L-glutamine (Sigma), 1% non-essential amino acid solution (Invitrogen), 1% penicillin-streptomycin (Invitrogen), and 0.1 mM β-mercaptoethanol (Sigma). The resulting DE cells were then differentiated in PE medium into PE cells for 6 d. PE medium is composed of DMEM-Glutamax (Invitrogen) supplemented with 0.5% B27 supplement (Invitrogen), 2 μM retinoic acid (Sigma), 2 μM dorsomorphin (A. G. Scientific), 10 μM SB431542 (Cayman), 5 ng/ml basic fibroblast growth factor (R&D Systems), and 250 nM SANT1 (Cayman). The resulting PE cells were incubated in EP medium for 4 d. EP medium is composed of DMEM-Glutamax containing 0.5% B27 supplement (Invitrogen), 50 μg/ml ascorbic acid (Sigma), 10 μM SB431542 (Cayman), 2 μM dorsomorphin (A. G. Scientific), and 10 μM DAPT (Cayman). The resulting EP cells were cultured in EC medium for 8 d. EC medium is composed of CMRL 1066 (Invitrogen) supplemented with 0.5% B27 supplement (Invitrogen), 25 mM glucose (Sigma), 0.5% penicillin-streptomycin (Invitrogen), 500 μM dibutyryl-cAMP (Santa Cruz Biotechnology, Santa Cruz, CA), 2 μM dorsomorphin (A. G. Scientific), 10 μM exendin-4 (Sigma), 10 μM SB431542 (Cayman), 10 mM nicotinamide (Sigma), and 50 μg/ml ascorbic acid (Sigma).

### Electroporation

Before electroporation, P-iKO cells were incubated in PE medium containing 1.2 μg/ml doxycycline (dox) for 3 d of PE differentiation to induce P-KO. On the final day in the PE stage, P-iKO PE cells were pre-incubated in PE medium containing 10 μM Y27632 for 1h and then dissociated. The cells were rinsed with DPBS and detached by treatment with Accutase (Innovative Cell Technologies, Inc., CA) at 37℃ for 10 min. Dissociated cells were centrifuged at room temperature (RT) at 300 x *g* for 3 min. After two washes in 1X Opti-MEM (Invitrogen), 1 × 10^6^ cells and 10 μg of DNA vector were mixed and electroporated at 150 V for 2.5 msec while suspended in 100 μl of Opti-MEM in a 2 mm-gap cuvette. Electroporated cells were cultured in mTeSR1 medium supplemented with 10 μM Y27632 at 37℃ and 5% CO_2_ for 1–2 d before western blot analysis.

### Western blot

Cultured cells were collected in a pre-cooled ep-tube using a cell scraper (SPL) and lysed in ice-cold RIPA buffer (InvivoGen) containing protease and phosphatase inhibitors. Lysates were sonicated briefly before any cellular debris was removed by centrifugation at 16,000 x *g* at 4℃ for 25 min. Total protein concentration was calculated via the Bradford protein assay (Bio-Rad, Hercules, CA). Approximately 10 μg of total protein was separated by SDS-PAGE electrophoresis and transferred to a nitrocellulose membrane with either 0.4 μm pores (GE Healthcare, Chicago, IL) or 0.2 μm pores (Invitrogen). After blocking with 5% BSA or 4% skim milk in TBST (10 mM Tris-HCl [pH 7.5], 150 nM NaCl, and 0.1% Tween 20), the membranes were incubated in 5% BSA or 4% skim milk TBST at 4℃ overnight with the appropriate primary antibodies. The primary antibodies used in this study were as follows: rabbit anti-PRMT1 (1:1000, Abcam, Cambridge, UK), mouse anti-NGN3 (1:500, DSHB, IA), mouse anti-FLAG (1:500, Santa Cruz), rabbit anti-AMDA (1:500, Cell Signaling Technology, Inc., Danvers, MA), rabbit anti-Methyl (mono) Arginine (1:500, ImmuneChem Pharmaceuticals Inc., British Columbia, Canada), HRP-conjugated rabbit anti-GAPDH (1:2000, Santa Cruz), rabbit anti-H4R3me2a (1:1000, CST), and rabbit anti-H3 (1:2000, CST). After three rinses with TBST, the membranes were incubated in 4% skim milk TBST at RT for 1 h with anti-mouse or anti-rabbit IgG HRP-linked secondary antibodies (CST). Protein bands were visualized in ECL solution (Merck KGaA, Darmstadt, Germany) using an ImageQuant LAS4000 (GE Healthcare). Band intensities were quantified using the ImageJ program (NIH, www.nih.gov).

### Quantitative RT-PCR

Total mRNAs were extracted from cultured cells using Easy-BLUE solution (Intron Biotechnology, Seongnam, Korea) and reverse-transcribed with a cDNA synthesis kit (BioAssay, Daejeon, Korea). Quantitative RT-PCR was carried out with SYBR Green and Taq DNA polymerase (SolGent Co., Ltd., Daejeon, Korea) on a Bio-Rad CFX96 Real-Time System (Bio-Rad). For analysis of relative expression, the Cq values for each gene were normalized with GAPDH mRNA levels using the ΔΔCq method. Primer sequences are listed in Table 1.

**Table 1.**
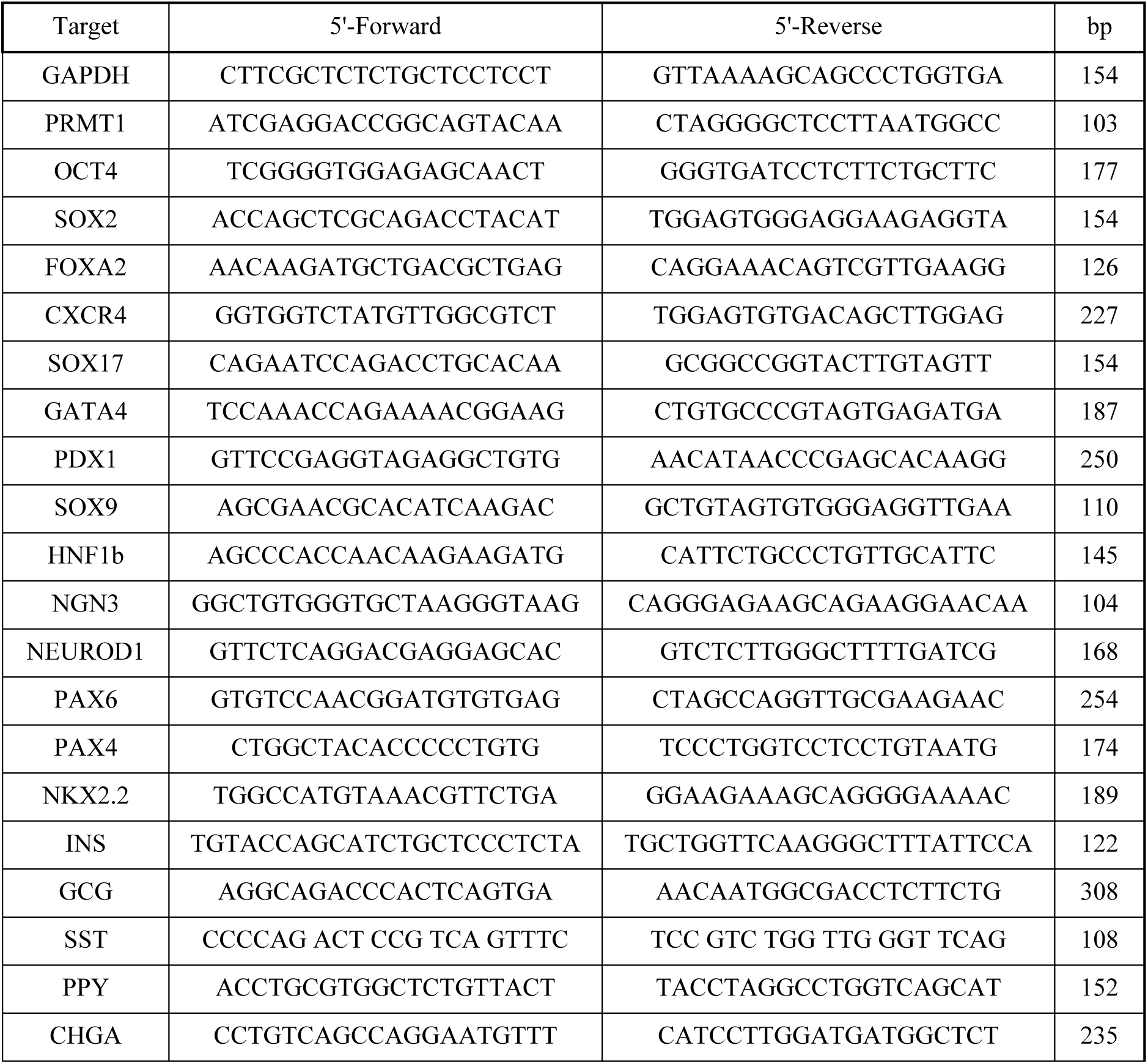
Quantitative RT-PCR primer list.

### Immunostaining

Cells were fixed with 4% formaldehyde solution (Sigma) at RT for 30 min. After three 5-min rinses in PBS, the cells were incubated in PBS containing 0.1% Triton-X (Sigma) for 30 min and blocked in PBS containing 1% BSA (Sigma) for 1 h. After the addition of the appropriate primary antibodies, the cells were incubated in blocking solution at 4℃ overnight. The primary antibodies were as follows: rabbit anti-SOX2 (1:200, CST), mouse anti-TRA 1–60 (1:200, Millipore, Billerica, MA), goat anti-OCT4 (1:500, Santa Cruz), mouse anti-TRA 1–81 (1:200, Millipore), rabbit anti-PRMT1 (1:1000, Abcam), rabbit anti-H4R3me2a (1:1000, CST), goat anti-SOX17 (1:400, R&D Systems), mouse anti-GATA4 (1:400, Santa Cruz), rabbit anti-PDX1 (1:1000, Abcam), goat anti-HNF1b (1:200, Santa Cruz), rabbit anti-HNF4a (1:400, Santa Cruz), mouse anti-NGN3 (1:200, DSHB), rabbit anti-SOX9 (1:2000, Abcam), guinea pig anti-insulin (INS, 1:1000, DAKO, Glostrup, Denmark), mouse anti-c-peptide (CPEP, 1:1000, Abcam), mouse anti-glucagon (GCG, 1:500, Sigma), mouse anti-pancreatic polypeptide (PPY, 1:200, R&D Systems), and rabbit anti-somatostatin (SST, 1: 1000, DAKO). After three washes in PBS, the cells were treated with Alexa-Flour 488- or 594-conjugated donkey antibodies (Abcam) at a dilution ratio of 1:1000 and further incubated in blocking solution at RT for 1 h. DAPI (Invitrogen) was used for nuclear counter-staining. After five washes in TBST, the stained cells were observed on either a fluorescence microscope (Olympus, Tokyo, Japan) or a confocal microscope (LSM780, Carl Zeiss, Oberkochen, Germany). Images were analyzed using the ZEN (Carl Zeiss) software and the ImageJ program.

### Immunoprecipitation

Transfected cells were sonicated in cold BC300 buffer containing 20 mM Tris (pH 7.9), 0.2 mM EDTA, 20% Glycerol, 300 mM KCl, 0.1% Tween 20, and 0.2 mM PMSF. After centrifugation at 16,000 x *g* at 4℃ for 25 min, the supernatants were aliquoted, 5% (v/v) for an input sample and 95% (v/v) for immunoprecipitation. The 95% (v/v) lysate aliquots were incubated with a monoclonal anti-FLAG M2 antibody (Sigma) at 4℃ overnight with mild agitation. Then, FLAG and M2 conjugates were washed three times in 700 μl of cold BC300 buffer. FLAG-NGN3 was eluted using FLAG peptide or 5X sample buffer. For co-immunoprecipitation of FLAG-NGN3 with HA-Ub, transfected HEK cells were treated with a proteasome inhibitor (20 μM MG132, Selleck Chemicals, TX) for 2 h. Eluted samples were boiled at 95℃ for 5 min and analyzed by western blotting.

### siRNA transfection

HEK cells were transfected with the appropriate siRNAs using the Lipofectamine RNAiMAX reagent (Invitrogen) according to the manufacturer’s instructions. This study used a human PRMT1-specific siRNA (cat#sc-41069, Santa Cruz) and a non-targeting control siRNA (Dharmacon, Lafayette, CO).

### Purification of recombinant proteins

To produce GST-tagged proteins, cDNAs encoding human NGN3 protein fragments were sub-cloned into pGEX4T-1 (GE Healthcare), expressed in *Escherichia coli*, and purified on Glutathione-Sepharose 4B beads (GE Healthcare) as previously described (Kim and Roeder, 2011). For FLAG-tagged proteins, cDNAs encoding mouse PRMT proteins were subcloned into pFASTBAC1 (Invitrogen) with a FLAG tag. Baculoviruses with individual Prmt cDNAs were generated according to the manufacturer’s instructions (Invitrogen). These were then used to infect Sf9 insect cells. The infected Sf9 cells were incubated in Grace’s insect medium (Invitrogen) supplemented with 10% FBS at 27℃ for 3 d. Finally, proteins were purified from the infected cells on M2 agarose (Sigma) as previously described (Kim and Roeder, 2011).

### In vitro methyltransferase assays

Purified substrates (i.e., 400 ng histone octamer or NGN3 proteins and 200 or 400 ng PRMT proteins) were mixed and then incubated in 20 μl of reaction buffer (25 mM HEPES at pH 7.6, 50 mM KCl, 5 mM MgCl_2_, and 4 mM DTT) supplemented with 200 μM cold SAM (*S*- adenosyl methionine) (NEB; for immunoblotting) or 1 µCi ^3^H-labelled SAM (PerkinElmer; for fluorography) at 37°C for 60 min, respectively. The proteins were resolved by 15% SDS-PAGE and subjected to immunoblotting (cold SAM) or autoradiography (radiolabeled SAM). Anti-dimethyl (asymmetrical) arginine (ICP0810) and anti-monomethyl arginine (ICP0801) antibodies were purchased from ImmuneChem.

### Luciferase reporter assays

After a 1-d incubation in DMEM containing 10% FBS, HEK cells were transfected with pCAG-FLAG-NGN3 WT- or R65A mutant-expressing vectors, the pGL3-NEUROD1 promoter (∼1 kb) firefly luciferase reporter, and a renilla luciferase-expressing vector at a 1:10:1 ratio using Lipofectamine 2000 (Invitrogen). The pCAG-FLAG backbone vector (empty) was used as a negative control. The transfected cells were cultured in the same medium for 2 d. Cell lysis and luciferase assays were performed using the Dual-Luciferase® Reporter Assay System kit (Promega Corporation, Madison, WI). Luciferase units were measured with a Spark microplate reader (Tekan Group Ltd., Männedorf, Switzerland). Firefly luciferase levels were normalized to renilla luciferase levels.

### NGN3 degradation assays

HEK cells were transfected with the pCAG-FLAG-NGN3 vector and then incubated in DMEM containing 10% FBS for 1 d. After changing to fresh medium supplemented with 10 μg/ml cycloheximide (CHX, Sigma), the transfected cells were immediately lysed with cold RIPA buffer containing fresh protease and phosphatase inhibitors at each CHX time point. Degradation of FLAG-NGN3 was analyzed by western blot.

### Statistical analysis

Statistical analyses of all data were carried out using GraphPad Prism 7 (GraphPad Software, Inc., La Jolla, CA). Each experiment was performed at least in triplicate. Data are presented as means ± SEM. Statistical significance was calculated via paired Student’s t-tests. P-values are expressed as * *p* < 0.05; ** *p* < 0.01; *** *p* < 0.001; **** *p* < 0.0001.

## Acknowledgements

We thank Prof. Yoon for providing ZFN-L, ZFN-R, pAAV-Neo_Cas9, and pAAV-Puro_siKO vectors. This study was supported by Korean Fund for Regenerative Medicine (KFRM) 21A0402L1-12, and the National Research Foundation (NRF) 2021R1A2C300510811, Republic of Korea.

## Supplemental Information

### Supplementary Figure Legends

**Figure S1.**
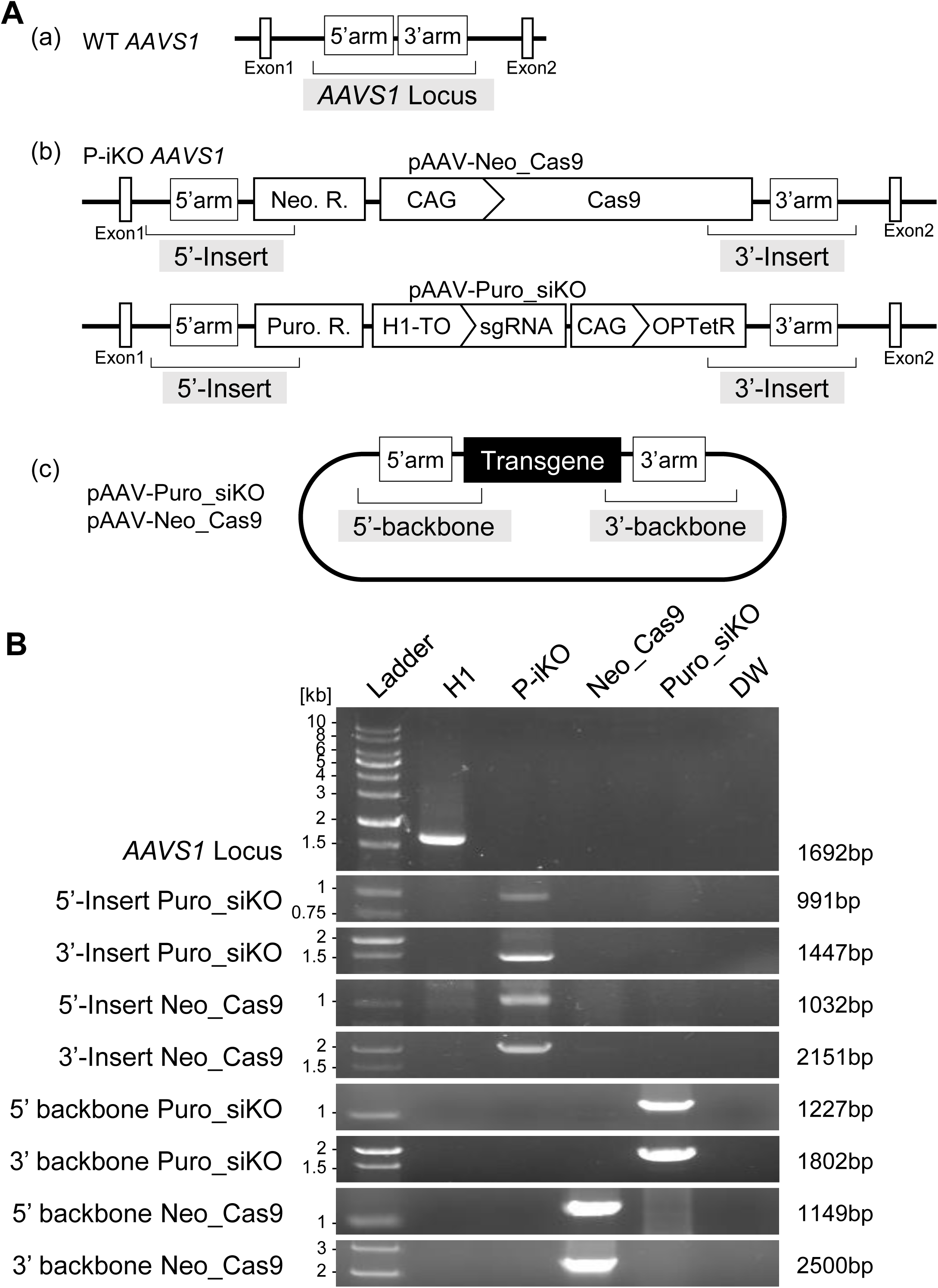
Genotyping of P-iKO hESCs. (A) Schematic P-iKO constructs for targeting the *AAVS1* genomic locus. *AAVS1* genomic locus of H1 hESCs (a), both P-iKO hESC targeted *AAVS1* alleles (b), and the *AAVS1* targeting vector (c). (B) Genotyping a P-iKO hESC line. PCR primers for genotyping are listed in ‘CRISPR Gene Editing’ (Snijders et al., 2019). DW was loaded as a negative PCR control.

**Figure S2.**
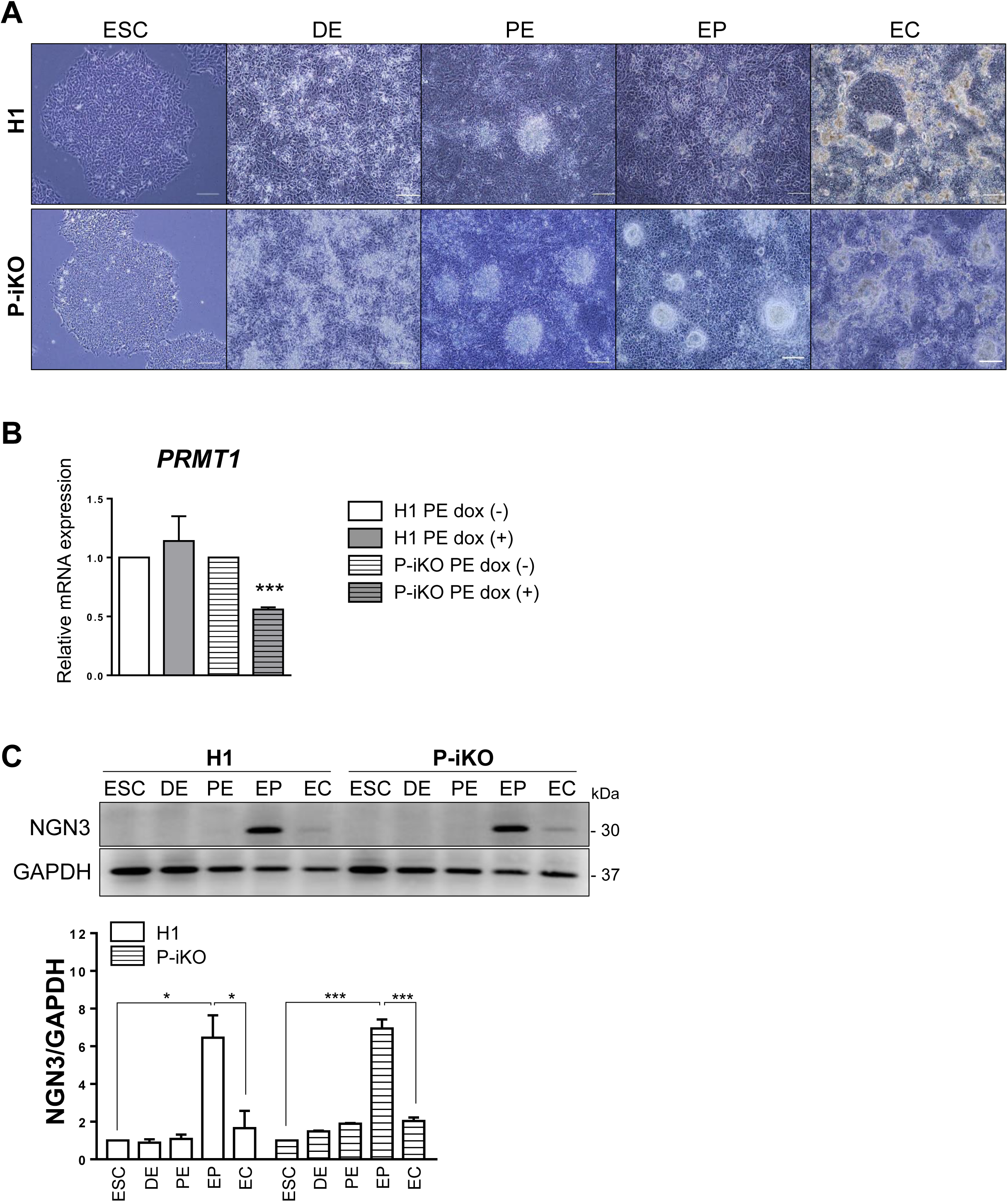
Differentiation of P-iKO hESCs into pancreatic ECs. (A) Cellular morphology at the various stages through which P-iKO hESCs progress as they differentiate into ECs. Scale bars, 200 μm. (B) Downregulation of *PRMT1* mRNA in P-KO PEs. Suppression of *PRMT1* was observed in dox (+) P-iKO PE cells on PE day 6 (PED6). \*\*\**p* < 0.001 (n = 4). (C) Expression of NGN3 in P-iKO hESCs as they progress through pancreatic EC development. Transient NGN3 expression was detected only in EP cells derived from both H1 and P-iKO hESCs. \**p* < 0.05, \*\*\**p* < 0.001 (n = 3).

**Figure S3.**
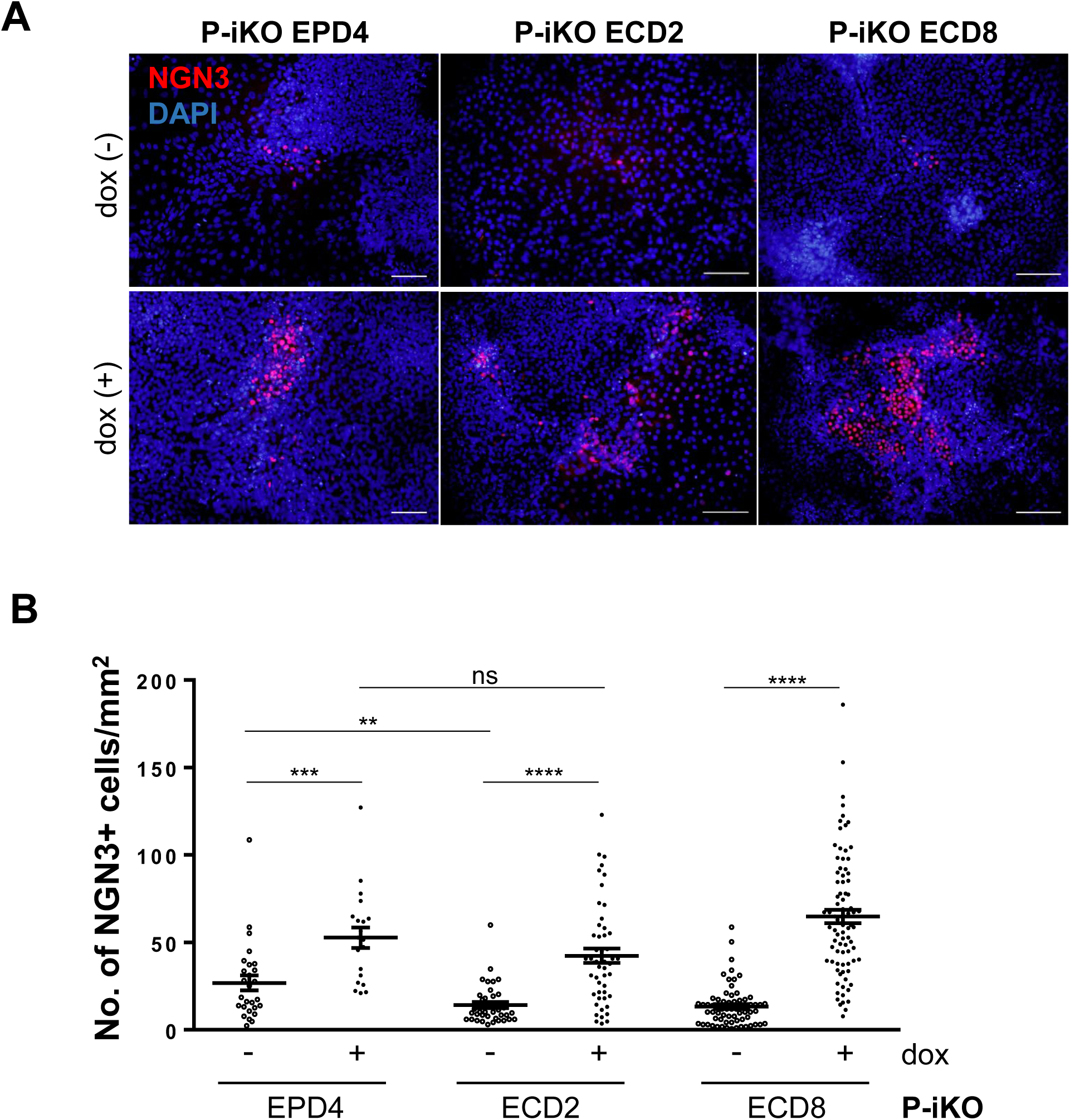
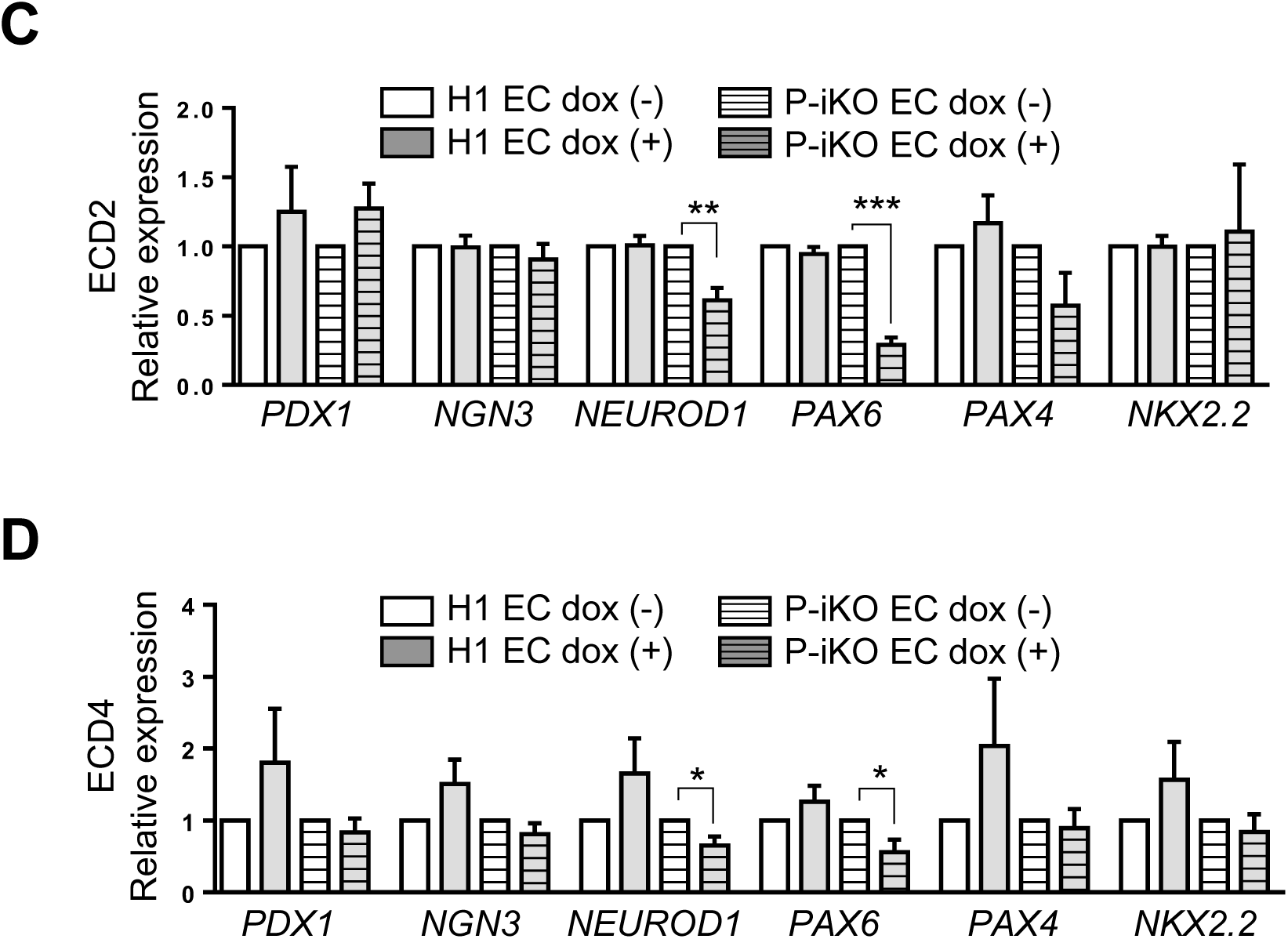
Effects of PRMT1 depletion in P-KO EPs and ECs. (A) More NGN3-positive cells among P-KO EPs and ECs on EP day 4 (EPD4), EC day 2 (ECD2), and EC day 8 (ECD8). Scale bars, 100 μm. (B) More NGN3-positive cells among P-KO EPs and ECs. \*\**p* < 0.01, \*\*\**p* < 0.001, \*\*\*\**p* < 0.0001 (n > 20). (C) Reduced mRNA expression of the NGN3 target genes *NEUROD1* and *PAX6* in P-KO ECs on EC day 2 (ECD2). \*\**p* < 0.01, \*\*\**p* < 0.001 (n = 4). (D) Significantly reduced mRNA expression of *NEUROD1* and *PAX6* in P-KO ECs on EC day 4 (ECD4). \**p* < 0.05 (n = 4).

**Figure S4.**
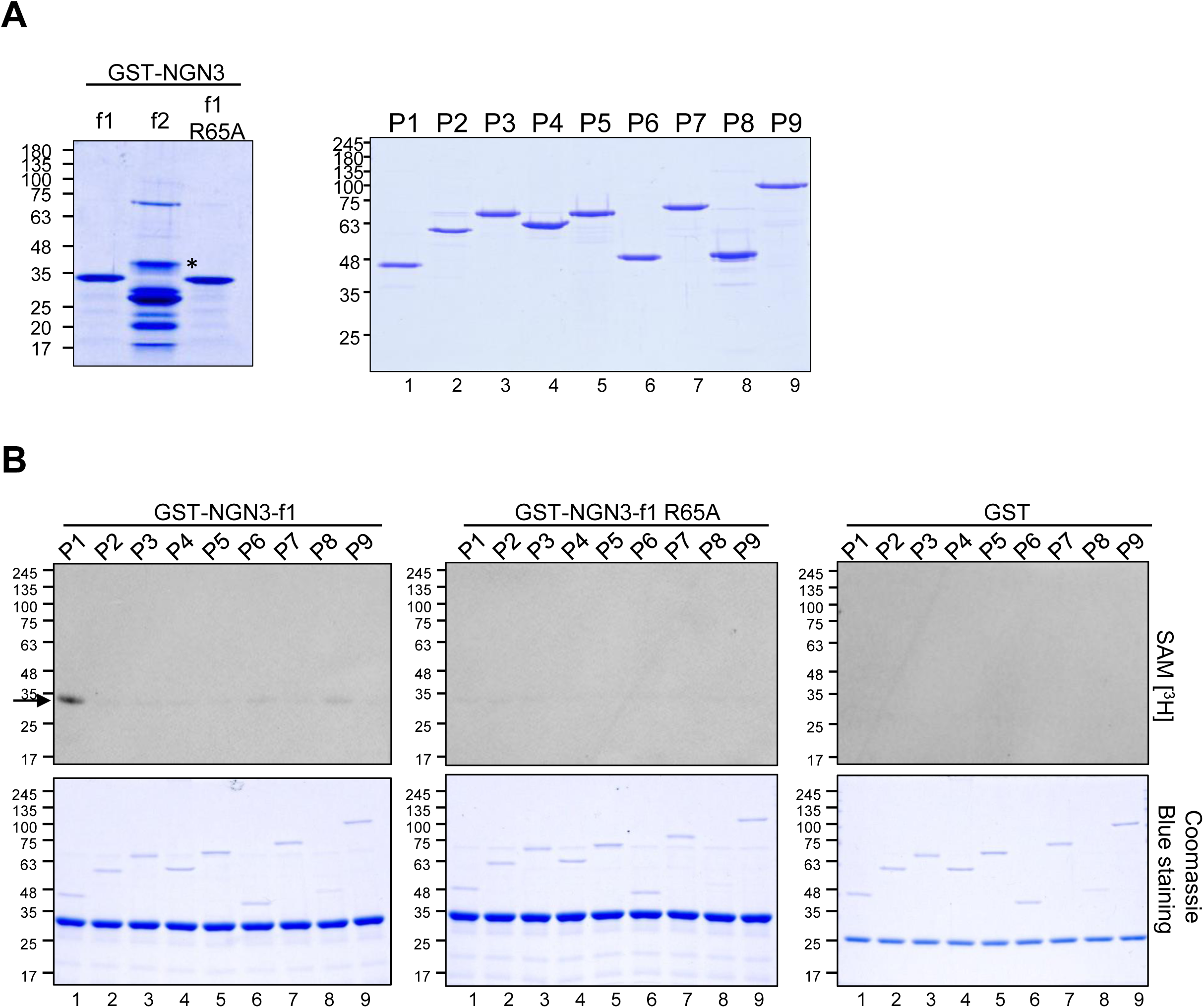
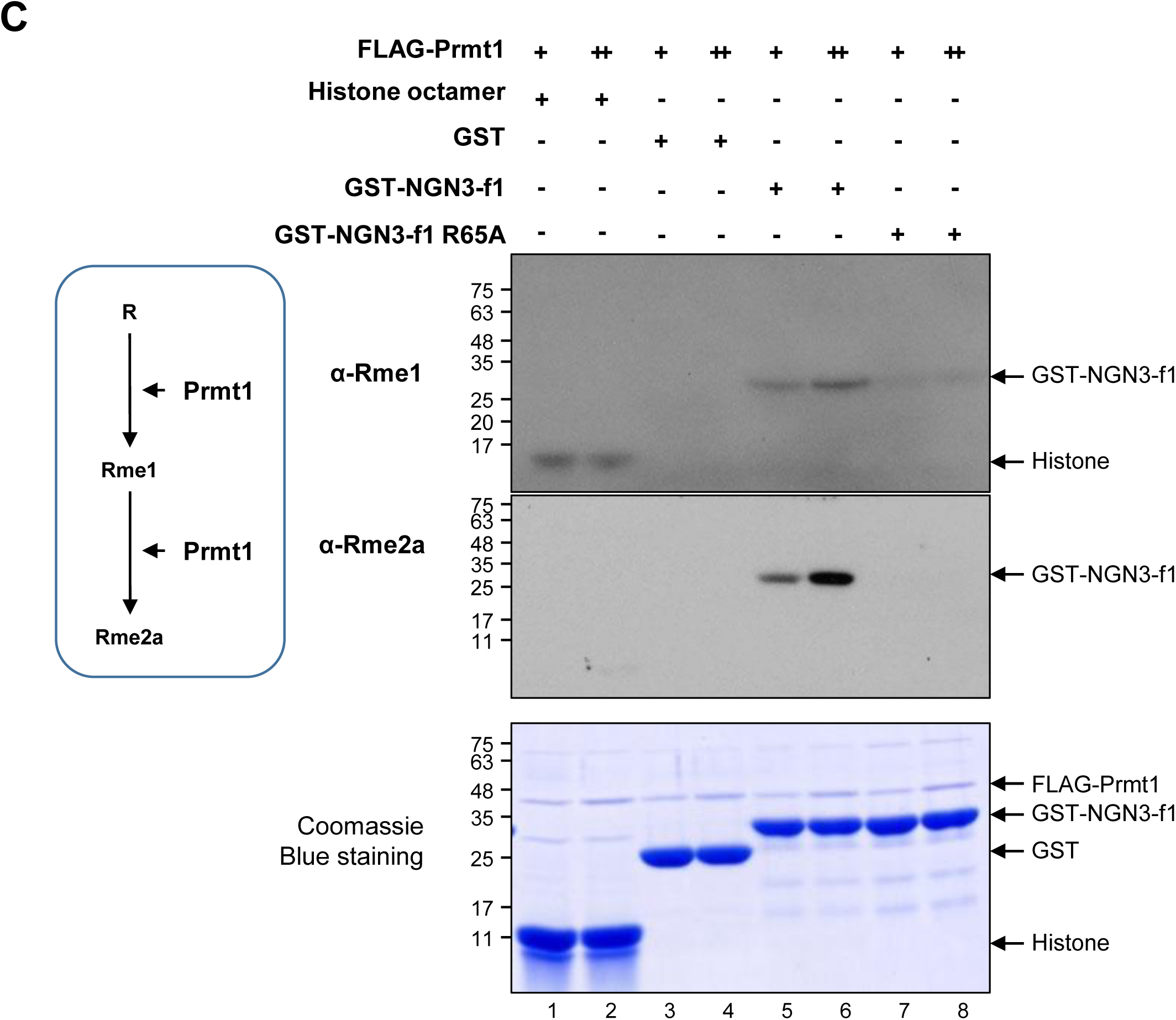
Arginine methylation of NGN3 by Prmt1. (A) Coomassie staining of GST-tagged NGN3 f1, f2 (asterisk), f1 R65A mutant (left), and Prmt family members (right). f, fragment of NGN3; P, Prmt. (B) Methylation of GST-tagged NGN3 f1 and f1 R65A mutant by Prmt family members. Only Prmt1 methylated an arginine residue in NGN3 f1 (arrow). (C) Methylation of GST-NGN3 f1 and R65A mutant by Prmt1. α-Rme1, mono-methylated arginine; α-Rme2a, asymmetrically di-methylated arginine. FLAG-Prmt1: +, 100 ng; ++, 200 ng. GST-NGN3 f1 and R65A: +, 400 ng, respectively.

**Figure S5.**
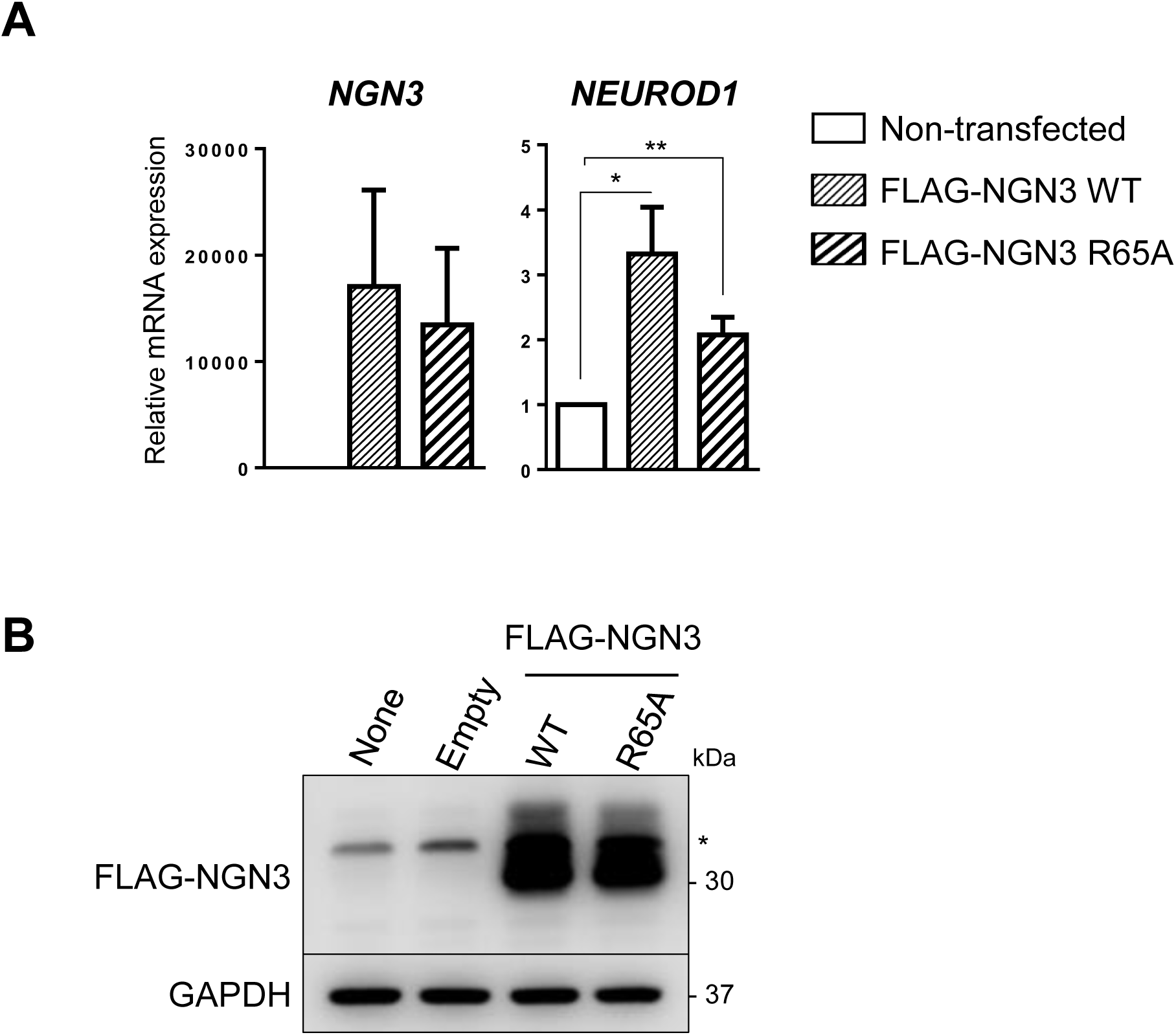
Expression of NGN3 and *NEUROD1*. (A) Relative mRNA expression of *NGN3* and *NEUROD1* in HEK cells transfected with pCAG-FLAG-NGN3 WT and R65A mutant. \**p* < 0.05, \*\**p* < 0.01 (n = 4). (B) Western blot analysis of pCAG-FLAG-NGN3-transfected HEK cells. “None” indicates cells incubated without any DNA vectors; “Empty” indicates cells transfected with pCAG-FLAG empty vector. The asterisk (*) indicates nonspecific bands.

**Figure S6.**
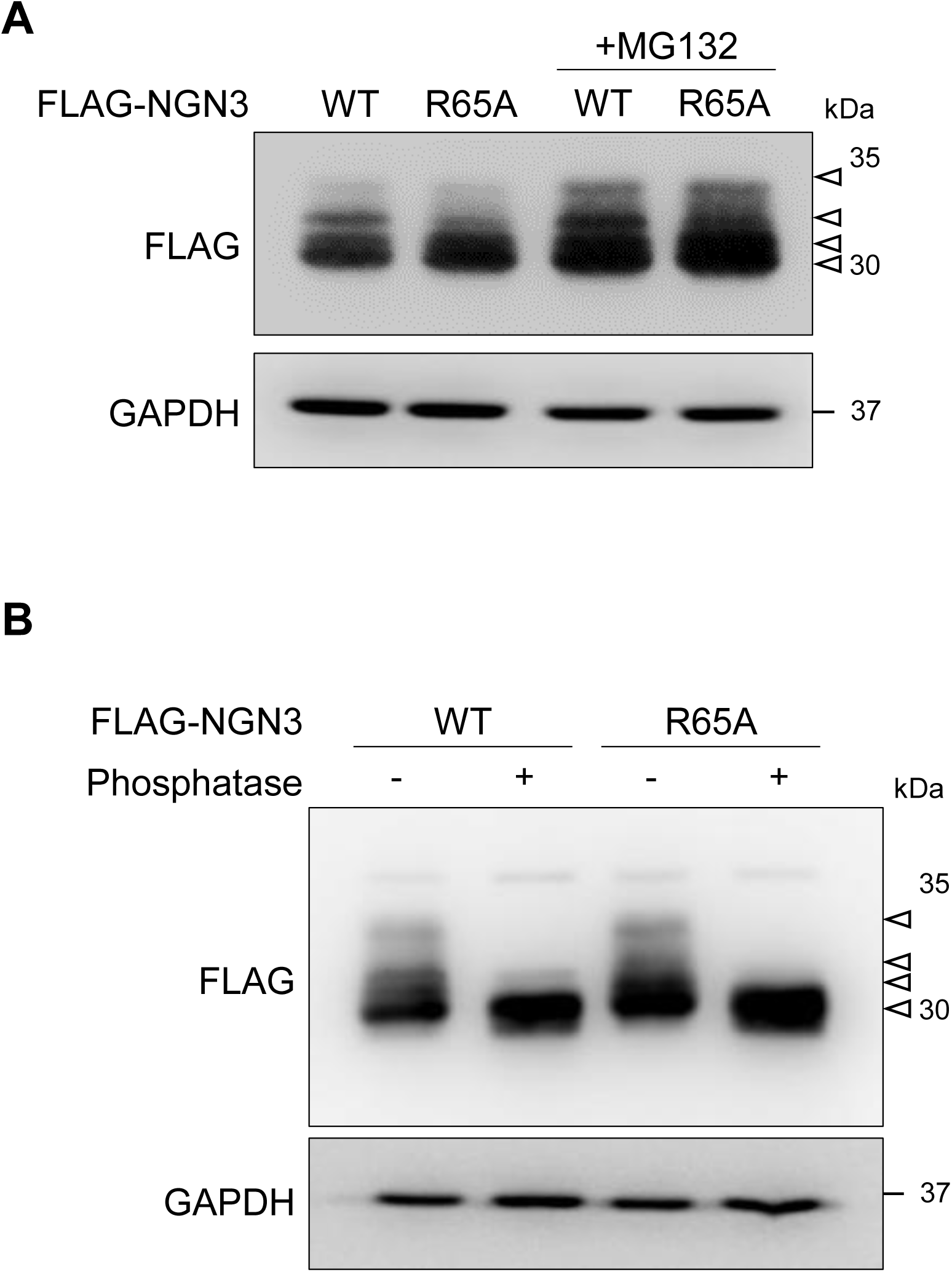
Phosphorylation of NGN3 WT and R65A mutant. (A) Western blot analysis of HEK cells transfected with pCAG-FLAG-NGN3 WT and R65A mutant. Multiple NGN3 bands (arrowheads) were detected for both the WT and R65A mutant. (B) Reduced NGN3 phosphorylation bands for both WT and R65A mutant induced by phosphatase treatment. Phosphorylated NGN3 bands (arrowheads) disappeared after phosphatase treatment (+).

